# Leveraging cell-type specificity and similarity improves single-cell eQTL fine-mapping

**DOI:** 10.1101/2025.03.05.641709

**Authors:** Chen Lin, Yingxin Lin, Wenxuan Li, Leqi Xu, Xiangyu Zhang, Hongyu Zhao

**Affiliations:** Department of Biostatistics, Yale University, New Haven, CT, USA 06510; Department of Computational Medicine, UCLA, Los Angeles, CA 90095

**Author notes:** To whom the correspondence should be addressed: Dr. Hongyu Zhao.

## Abstract

Identifying cell-type-specific expression quantitative trait loci (eQTLs) is important to understanding the genetic regulation of gene expressions at the cell-type level and its relevance to complex traits. However, existing eQTL fine-mapping methods are limited in power and accuracy when cell types are analyzed separately. To improve eQTL mapping, we present CASE, a Bayesian framework to perform **C**ell-type-specific **A**nd **S**hared **E**QTL fine-mapping that simultaneously analyzes multiple cell types. CASE can effectively capture effect-sharing patterns across cell types while disentangling the confounding effects of linkage disequilibrium (LD). We demonstrate that CASE outperforms the existing single-trait (SuSiE) and multi-trait (mvSuSiE) eQTL methods through comprehensive simulations. When applied to the OneK1K data, CASE identified more genetic regulations of gene expressions, better capturing cell type specificity and functionally enriched and disease-associated eQTLs. The CASE framework for single-cell eQTL fine-mapping can be broadly applied to multi-tissue and multi-trait genetic studies.

## Introduction

Expression quantitative trait locus (eQTL) analysis aims to identify genetic variants, such as single-nucleotide polymorphisms (SNPs), associated with gene expression levels. Complementing genome-wide association studies (GWAS), which primarily link genetic variants to complex traits, eQTL analysis offers insights into how genetic variants affect disease risk by regulating gene expressions^1-3^. However, two main challenges remain in current eQTL analysis. The first challenge is linkage disequilibrium (LD) among SNPs, resulting in a block of highly correlated SNPs near the causal eQTL(s), all displaying significant p-values. Statistical fine-mapping addresses this challenge by disentangling LD and distinguishing putative causal variants from merely correlated ones^4,5^. Fine-mapping results therefore may more effectively implicate biological pathways driving disease onset and identify novel biomarkers. While most fine-mapping efforts have focused on GWAS, recent studies, such as the GTEx v10 release, have extended fine-mapping methods to eQTL data.

The second challenge arises from the fact that most existing eQTL studies analyze bulk tissues, capturing only average genetic regulation across cell types. This approach overlooks cell-type-specific or context-dependent eQTLs^6,7^, potentially contributing to the limited overlap between GWAS and eQTL findings. As a result, despite the success of eQTL studies in interpreting GWAS signals, only a small proportion of disease-associated loci can be co-localized to eQTLs^8,9^, and known eQTLs only explain a small fraction of heritability for complex traits^10-12^. The recent development of single-cell technologies and the availability of single-cell RNA sequencing (scRNA-seq) data from genotyped individuals enable the identification of cell-type-specific eQTLs^13,14^, leading to insights on genetic regulation at the cellular level and its relevance for complex diseases and traits.

Considering these challenges, integrating scRNA-seq data with fine-mapping methods can help understand genetic regulations of gene expressions. However, most fine-mapping methods analyze cell types separately, ignoring the shared regulatory patterns across cell types, as validated by recent studies^13,15^. Consequently, separate analyses of individual cell types do not benefit from shared regulatory patterns across related cell types. In contrast, previous multi-tissue analyses in bulk eQTL data have shown that jointly modeling tissues enhances power^16-18^, suggesting that similar approaches could benefit single-cell eQTL analysis. Currently, no study has comprehensively addressed the simultaneous disentanglement of LD and joint modeling of effect-sharing patterns across cell types in single-cell eQTL analysis.

One potential approach to addressing these challenges is to utilize multi-trait fine-mapping methods by considering each cell type as a separate trait. While some multi-trait fine-mapping methods were proposed to address this issue (as summarized in Zou *et al*.^19^), most of these methods (e.g. flashfm, PAINTOR) are limited to two or three traits due to modeling or computational constraints^20-23^, making them unsuitable for single-cell eQTL data with many cell types involved. Other methods, such as CAFEH^24^ and HyPrColoc^25^, do not account for the effect-sharing patterns and correlations across traits. Among the available methods, only mvSuSiE models effect-sharing patterns for multiple traits, which first applies a multi-trait marginal testing method, mashr, to learn the patterns of shared genetic effects, and subsequently applies SuSiE for fine-mapping.

In mvSuSiE, there are two approaches for modelling effect-sharing patterns: the data-driven method and the canonical method. However, both have limitations that impact their ability to control false discoveries and maintain statistical power. The canonical approach uses pre-fixed correlation matrices representing different sharing patterns, which may not align with the true correlation structure. The data-driven approach relies on mashr to estimate sharing patterns from eQTL summary statistics. However, mashr, designed for marginal association analyses, does not account for local LD structures among SNPs, leading to two major issues. First, correlations estimated across cell types are conflated with LD, failing to reflect true correlations of the underlying eQTL effect sizes, potentially resulting in false positives and negatives (See examples in Fig. 1b). Second, these patterns are averaged across genes without accounting for variations in local genetic regions, where average patterns may not represent a specific gene accurately.

**Fig. 1.**
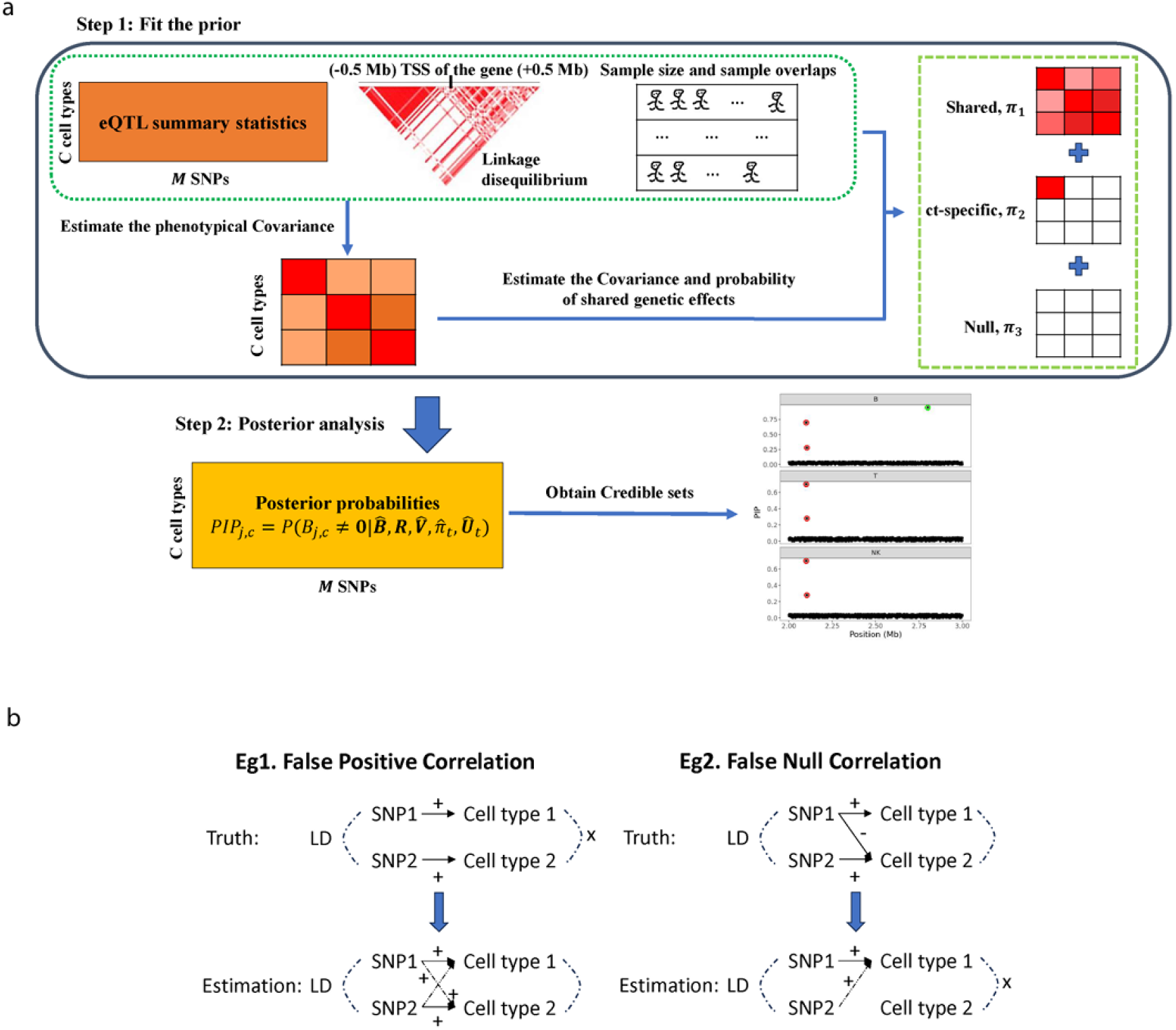
Schema of CASE. **a**, The CASE workflow. The input data consist of eQTL summary statistics, LD matrix, sample size and sample overlap information. The phenotypical and effect-sharing covariances are estimated from the input data. The sketch shows a simple example of three patterns of effect sharing across three cell types, while CASE can fit a wide range of patterns for multiple cell types based on the input. The fitted prior is then used to obtain the posterior probabilities and credible sets per cell type. **b**, Two illustrations of false positives and negatives introduced when effect-sharing patterns are estimated mixing with LD. In example one, two cell types are independent while the causal eQTLs are in LD, which may lead to a false positive correlation between cell types. In example two, eQTLs in LD have opposite effect sizes, which cancel out in cell type 2, leading to false uncorrelated estimates between cell types.

More importantly, the covariance structures from both approaches lack flexibility regarding the number of shared cell types. The covariance matrices assume effects that are either unique to one cell type or shared across all cell types, with diagonal elements either all or only one to be nonzero. This approach only captures extreme cases of sharing patterns—either cell-type-specific or universally shared—without allowing for configurations where genetic regulation is shared across a subset of cell types. This may lead to inflated false discoveries by overemphasizing the two extremes of cell-type sharing. In addition, single-cell data commonly have missing samples for some cell types due to the limited sequencing depth, while mvSuSiE cannot accommodate the missing samples.

In this paper, we propose a novel empirical Bayesian model named CASE, which more appropriately leverages cell-type specificity and similarity to improve statistical power in eQTL fine-mapping. The main advantage of CASE lies in its flexible approach to learning a broad range of effect-sharing patterns from eQTL summary statistics while accounting for LD, allowing for variable numbers of shared cell types and diverse correlation structures to address limitations in previous methods. It also accommodates missing samples across cell types. We demonstrate the efficacy of CASE through simulations and application to the OneK1K dataset^13^, highlighting its capacity to identify functionally enriched and disease-relevant eQTLs by offering a robust approach for modeling cell-type specificity and similarity in fine-mapping studies.

## Results

### Overview of CASE

The CASE framework employs an empirical Bayesian approach consisting of two main steps: prior fitting and posterior analysis (Fig. 1a). The input of CASE requires eQTL summary statistics (e.g., z-scores or marginal effect sizes with standard errors) from each cell type, a reference LD matrix, sample sizes per cell type, and information on sample overlaps between cell types. In the prior fitting step, it estimates the phenotypic covariances due to sample overlaps and the eQTL effect-sharing patterns across cell types. In the posterior analysis step, it calculates the posterior probability of each SNP being a true eQTL for each cell type, and aggregates these results to identify putative causal eQTL sets. To account for the varied local LD patterns across genomic regions, CASE analyzes each gene individually, focusing on the *cis*-regulatory region (500 Kb upstream and downstream of the transcription start site, TSS).

#### CASE Model

Suppose there are *C* cell types, each with sample size *N*_*C*_. For a given gene with *M* SNPs in its *cis*-region, the genetic regulation of gene expression levels across cell types can be formulated as the following multivariate multiple regression model:

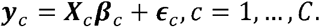

Here, 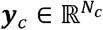 denotes the standardized gene expressions in cell type *C* after appropriate normalization and covariate adjustments (e.g., sex, age, genotype principal components (PCs), and gene expression PCs), 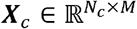 denotes the standardized genotypes within the *cis*-region of the gene, **β**_*C*_ ∈ ℝ^*M*^ represents the *cis*-eQTL effect size vector, and 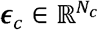 is the noise vector.

We assume that the noise terms are independent across individuals but correlated for the overlapped individuals across cell types, which results in a phenotypic covariance matrix ***V***_*y*_. We denote the matrix of *cis*-eQTL effect sizes as *B* = (***β***_1_, …, ***β***_*C*_) ∈ ℝ^*M*×*C*^, and assume that each row *B*_*j*_·, representing the effect size of the SNP *j* across cell types, follows a mixture normal distribution with T components:

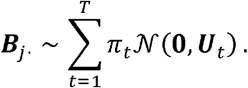

Here, 𝒩(·) denotes a multivariate normal distribution, each covariance matrix ***U***_*t*_ ∈ ℝ ^*C*×*C*^ characterizes a specific pattern of shared effect sizes, and π*t* indicates the probability of each pattern. For example, if no effect exists in cell type *C* under a pattern ***U***_*t*_, the *C*th row and column of ***U***_*t*_ contain only zeros. The number of patterns, *T*, is determined based on the observed patterns in eQTL summary statistics (Methods). The last pattern ***U***_*T*_ is fixed zero, representing SNPs with no eQTL effect in any cell type. We further assume independence among the *cis*-eQTL effect size, noise and genotype. These assumptions are commonly used in models for GWAS and eQTL studies^18,19^.

Let 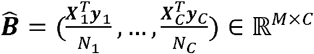 denote the eQTL summary statistics from gene-SNP marginal association tests. Given the true *cis*-eQTL effect size, 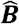 asymptotically follows a matrix normal distribution (Methods):

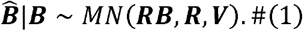

Here, *MN*(·) denotes a matrix normal distribution, V is the sample-adjusted covariance of gene expression, 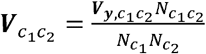, where 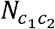 denotes the overlapped sample size between cell types *c*_l_ and *c*_2_, and ***R*** ∈ ℝ^*M*×*M*^ represents the LD matrix, which can be estimated from the study population or reference panels. To note, Equation (1) aligns with the well-studied single-trait (*C* = 1) models^24-26^, as the diagonal elements are 1/*Nc* in ***V*** due to the standardized gene expressions.

#### Step1: Prior Fitting

In the prior fitting step, we first estimate the sample-adjusted phenotypic covariances, V, using weak eQTL signals similar to mashr. Then, we apply a Monte Carlo Maximization-Expectation (MCEM) algorithm^27-29^ to estimate the sharing patterns, ***U***_*t*_, and their weights, π_*t*_ (Methods). In brief, this process treats the true effect sizes and pattern indicators as missing variables, iteratively maximizing the expected likelihood of the effect sizes given the summary statistics^30^. Because the SNPs in the same LD block exhibit dependencies in the effect-sharing patterns, the expectation of the likelihood can be extremely complicated. We propose to use a Markov chain Monte Carlo (MCMC) algorithm to approximate the distributions instead of using an explicit form.

#### Step2: Posterior Inference

For posterior inference, we obtain the posterior inclusion probability 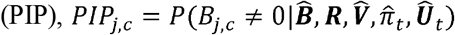 for each SNP *j* in cell type *C*, through the same MCMC algorithm, with multiple runs with different random initializations to ensure robust posterior estimates^31,32^. Finally, the posterior results are aggregated into credible sets^5^ based on LD structures, creating the smallest subsets of correlated variants containing at least one causal SNP (Methods).

### CASE consistently outperforms the existing methods in identifying true causal eQTLs in simulation studies

We conducted simulation studies to compare CASE with mvSuSiE, a multi-trait fine-mapping method, and SuSiE, a single-trait method. The first scenario evaluated the overall performance in identifying causal eQTLs under complex sharing patterns across varying sample sizes and heritability levels. The second scenario focused on Type I error control. For each simulation, we generated gene expression data with 1–5 true eQTLs across five cell types (Methods). The cis-heritability decreased across cell types, with cell type 1 having the highest and cell type 5 having the lowest heritability. This design allowed us to assess how CASE leverages highly heritable cell types to improve power in lowly heritable ones.

The first simulation scenario reflected real genomic complexity. Each eQTL was randomly classified as cell-type-specific or shared among a random number of cell types with equal probability. To understand the impacts of the sample size and heritability level, we varied the number of individuals and multiplied the *cis*-heritability with different multipliers (Methods).

We evaluated the performance using four metrics: (1) power, the proportion of true eQTLs found within credible sets, (2) coverage, the proportion of credible sets containing at least one of the true eQTLs, analogous to one minus false discovery rate (FDR) regarding credible sets, (3) size, the average number of variants per credible set, and (4) purity, the average minimum correlation among variants within credible sets. Power and coverage assess the capability of a method to detect true signals and control false discoveries, while size and purity evaluate its ability to distinguish causal SNPs from correlated ones.

As shown in Figs. 2a–b, CASE consistently outperforms mvSuSiE and SuSiE in identifying true causal eQTLs across cell types, improving power with increasing sample size and heritability. The power of SuSiE increases with *cis*-heritability, while CASE largely reduces the power discrepancies across cell types by leveraging cross-cell-type information. As for coverage, mvSuSiE fails to achieve the target of 95% in all cases, while CASE consistently has higher coverage than mvSuSiE, meeting the target in most scenarios. Although SuSiE has the highest coverage, it exhibits overly stringent FDR control, resulting in substantial power loss. Interestingly, unlike power, there is little discrepancy in coverage across cell types with different heritability values. Both CASE and mvSuSiE demonstrate comparable size and purity metrics, which are significantly better than those obtained by SuSiE (Supplementary Fig. 1). The results also suggest that a larger sample size and higher *cis*-heritability lead to improved power, coverage, size and purity for all methods.

**Fig. 2.**
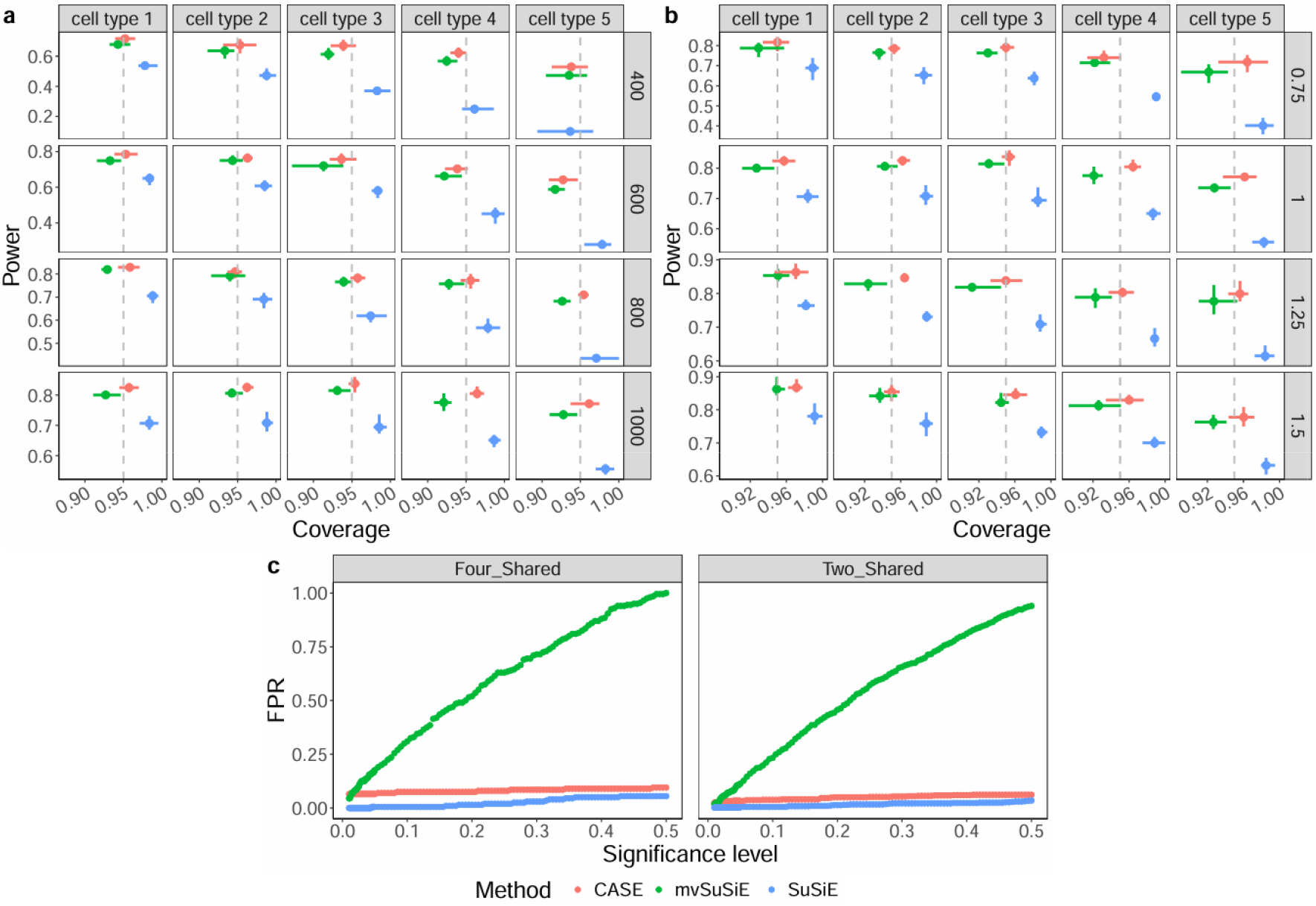
Simulation results. We compare CASE (red), mvSuSiE (green), and SuSiE (blue) in simulation studies. **a-b**, Power and coverage for varied sample size (**a**) and varied heritability multiplier (**b**). Each block represents a cell type indexed from one to five in a simulation setting. Error bars indicate upper 90% and lower 10% percentiles across 100 simulations. The grey vertical lines represent the 95% coverage target. **c**, False positive rates (FPR) in scenarios focusing on over-sharing error. The significance level was varied from 0 to 0.5, corresponding to the coverage threshold varying from 1 to 0.5 when aggregating SNPs into credible sets. The left plot shows the scenario where eQTL effects are shared across four cell types except for the remaining one, and the right plot shows the scenario where eQTL effects are only shared across two cell types and no eQTLs in the other three cell types.

### CASE better controls false positive for “over-sharing error”

While leveraging shared patterns across cell types enhances power, improperly weighting on these patterns can lead to misidentified eQTLs. For instance, if a SNP affects a gene in all immune cell types except for the B cells, multi-trait methods might incorrectly mark it as an eQTL for B cells due to over-emphasis on the shard-by-all patterns. We refer to this as an “over-sharing error”, a statistical artifact not commonly seen in single-trait methods.

In the second simulation, we assessed the control of over-sharing errors by CASE, mvSuSiE, and SuSiE. The simulation setup mirrored the first scenario, designed with two cases: (1) cell type 5 was assigned zero cis-heritability to represent a “null” cell type, assessing over-sharing error caused by shared-by-all patterns, and (2) cell types 3 to 5 were assigned zero heritability, assessing over-sharing error caused by shared-by-some patterns (Methods). To examine the sensitivity to the number of shared cell types in true eQTL effect sizes, we generated credible sets at varying significance levels.

The results (Fig. 2c) show that SuSiE yields almost no false positives as expected. CASE also effectively controls the false positive rates well, demonstrating robust performance. In contrast, mvSuSiE exhibits inflated false positive rates regardless of the number of truly shared cell types. This issue arises because the data-driven and canonical priors of mvSuSiE only characterize shared-by-all and cell-type-specific patterns. When only a subset of cell types shares strong eQTL effects, mvSuSiE will derive low posterior probabilities of the cell-type-specific patterns, resulting in the over-weighting of the shared-by-all patterns.

Overall, these two simulations demonstrate superior flexibility and robustness of CASE in characterizing diverse effect-sharing patterns, accurately identifying underlying eQTLs and effectively controlling false discovery rates, particularly for over-sharing errors, compared to mvSuSiE.

### CASE identifies more eGenes with more appropriate sharing patterns in the OneK1K data

To evaluate the performance of CASE in real data analysis, we analyzed the OneK1K dataset^13^, which contains 1.27 million peripheral blood mononuclear cells (PMBCs) collected from 981 genotyped healthy European donors. Among the 11,704 candidate genes, CASE identifies 5,043 eGenes with at least one eQTL in at least one cell type, representing a 12.6% increase over mvSuSiE and a 32.5% increase over SuSiE. Most eGenes are identified by all three methods (Fig. 3a). Here, we use “eQTLs” for putative causal SNPs within credible sets, rather than the associated ones. CASE-identified eGenes are further compared with the Genotype-Tissue Expression (GTEx) bulk eQTLs and fine-mapping results from whole blood (v10). On average, 83.2% eGenes are validated through GTEx studies, while only 68.9% eGenes are overlapped with GTEx fine-mapping results (Supplementary Fig. 2). Compared to the bulk-tissue results, the novel findings in single-cell results underscore the potential of CASE in uncovering novel regulations of gene expressions at the cell-type level.

**Fig. 3.**
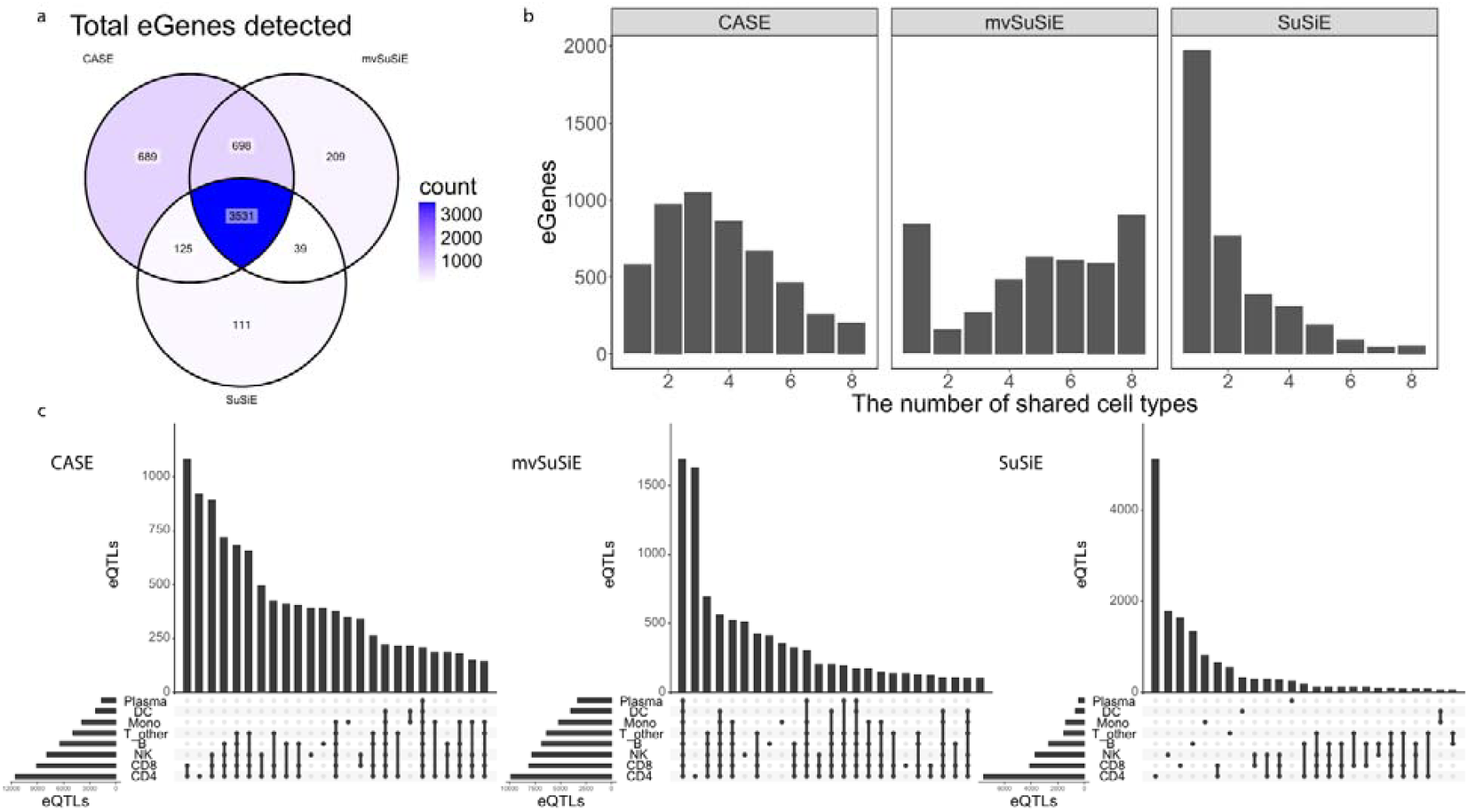
Fine-mapping results in the OneK1K eQTL data. **a**, Venn diagram of the number of eGenes (with at least one credible set) identified by CASE, mvSuSiE, and SuSiE. **b**, The number of eGenes shared across different numbers of cell types for CASE, mvSuSiE, and SuSiE. **c**, The number of eQTLs in the top 20 effect-sharing patterns for eight cell types identified by CASE, mvSuSiE, and SuSiE, respectively.

We further investigated the shared patterns of eGenes and eQTL across cell types (Fig. 3b-c). We expect similar cell types will share more eGenes while diverse cell types share less, leading to a variable number of shared cell types and various sharing patterns. In terms of the number of shared cell types for eGenes (Fig. 3b), SuSiE predominantly captures cell-type-specific patterns but misses numerous shared effects, whereas mvSuSiE excessively weights on both cell-type-specific and shared-by-all patterns, resulting in an unexpected trend where the number of eGenes increases with the number of shared cell types. This suggests potential false positives from over-sharing errors, as also evidenced in simulation studies. In contrast, CASE shows a more balanced sharing trend, accommodating variable numbers of shared cell types appropriately. The top 20 sharing patterns of eQTLs (Fig. 3c) reveal that the total eQTL counts were correlated to the cell type proportions (Supplementary Table 1), while both mvSuSiE and CASE improve the power especially for those less abundant cell types. Consistent with the eGene findings, SuSiE lacks power for shared effects, while mvSuSiE over-emphasizes the universal sharing. In contrast, CASE effectively captures a range of sharing patterns, highlighting that leveraging information from larger cell types (e.g., CD4 positive T cells) can enhance eQTL fine-mapping in smaller but related cell types (e.g., CD8 positive T cells and other T cells).

### CASE characterizes cell-type specificity and similarity

The average eQTL effect-sharing pattern estimated by CASE align with cell differentiation patterns and clustering (Fig. 4a). Similar cell types, such as T cell subtypes, including CD4 positive T cells (CD4), CD8 positive T cells (CD8), and other T cells (T_other), and dendritic cells (DC) and monocytes (Mono), display higher rates of shared eQTL effects compared to more distinct cell types. Most shared effect sizes exhibit consistent directional signs, contributing to a positive correlation among immune cell types, likely reflecting their similar regulatory functions. Notably, plasma blast cells (Plasma) display fewer shared effects with other cell types, including closely related B cells, possibly due to their low abundance and small sample size, which reduced the power of detecting causal eQTLs even with multi-trait approaches.

**Fig. 4.**
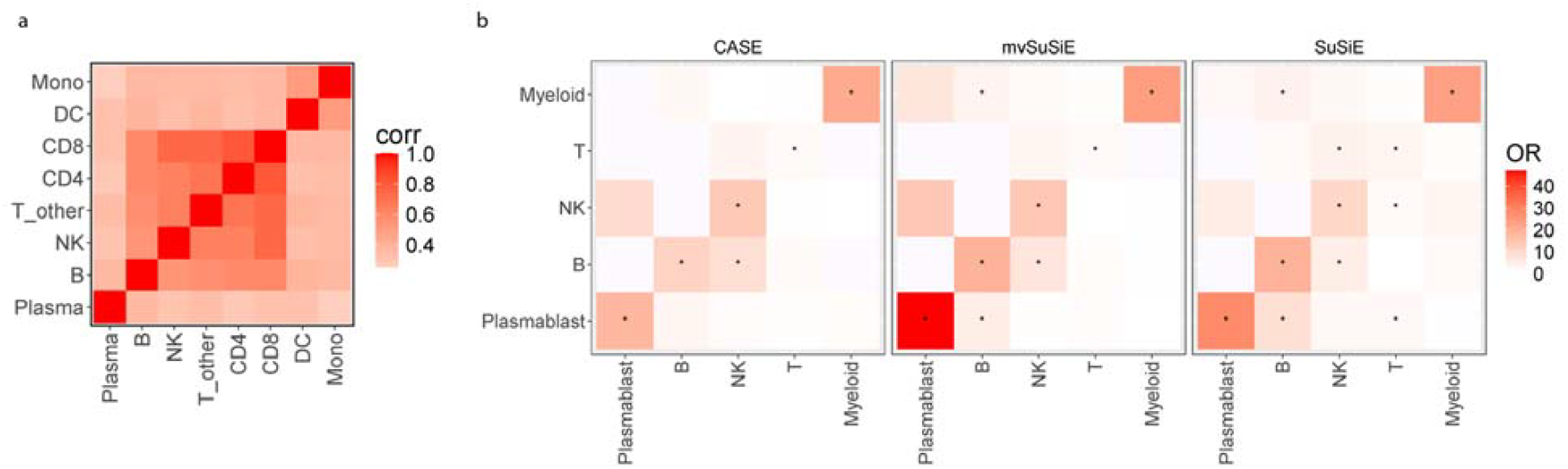
Cell-type specificity and similarity in OneK1K data. **a**, Heatmap of the average correlation of the fitted eQTL effects across all tested genes. **b**, Heatmaps showing the enrichment of cell-type-specific eGenes identified by three methods against cell type markers for each cell type. Colors represent odds ratio values, and stars indicate significance based on Fisher’s exact test with an adjusted p-value threshold of less than 0.05. T: aggregating all T subtypes, including CD4, CD8 and other T cells. Myeloid: aggregating DC and Mono.

We further examined the specificity of the cell-type-specific eGenes identified by three methods, hypothesizing that eGenes unique to certain cell types are more likely to be the markers for those cell types. To test this, we performed an enrichment analysis of cell-type-specific eGenes against marker genes of those cell types (Method). The results show that cell-type-specific eGenes are significantly enriched for the markers of their corresponding cell types across all three methods (Fig. 4b). However, mvSuSiE and SuSiE display more enrichments in unmatched cell types compared to CASE. This indicates that the cell-type-specific eGenes identified by CASE exhibit a stronger and more consistent enrichment within the appropriate cell-type-specific contexts, suggesting greater accuracy in identifying cellular-context relevant eGenes.

### CASE identifies functionally enriched eQTLs in disease-related cell types

To demonstrate the potential of CASE to detect signals while reducing false positives, we analyzed the putative causal variants of two genes and their regulatory mechanisms. The first gene, IRF5, is a protein-coding gene well-known for its impacts on autoimmune diseases such as systemic lupus erythematosus (SLE)^33^ and rheumatoid arthritis (RA)^34^, with particular relevance in myeloid cells and B cells. All three methods identify credible sets in DC and Mono (Figs. 5a-b). These credible sets contain two common variants, rs6969930, located in a region of open chromatin reflected by the DNase I enzyme (DHS), and rs3823536, which functions as both a promotor and a super-enhancer at transcription start sites. Additionally, rs3823536 has been validated in GTEx bulk eQTL fine-mapping of whole blood tissue, suggesting its potential causal role. Both CASE and mvSuSiE identify credible sets in additional cell types, including B cells, evidenced by nominally significant p-values. In contrast, SuSiE lacked the power to identify signals in these cell types. Interestingly, mvSuSiE identifies credible sets with the same two lead variants among all cell types; however, SNPs within these sets show non-significant marginal p-values in certain cell types that CASE does not identify, as exampled by Plasma (Fig. 5a). This suggests that mvSuSiE may be more susceptible to false positives due to its tendency to over-emphasize the fully shared patterns.

**Fig. 5.**
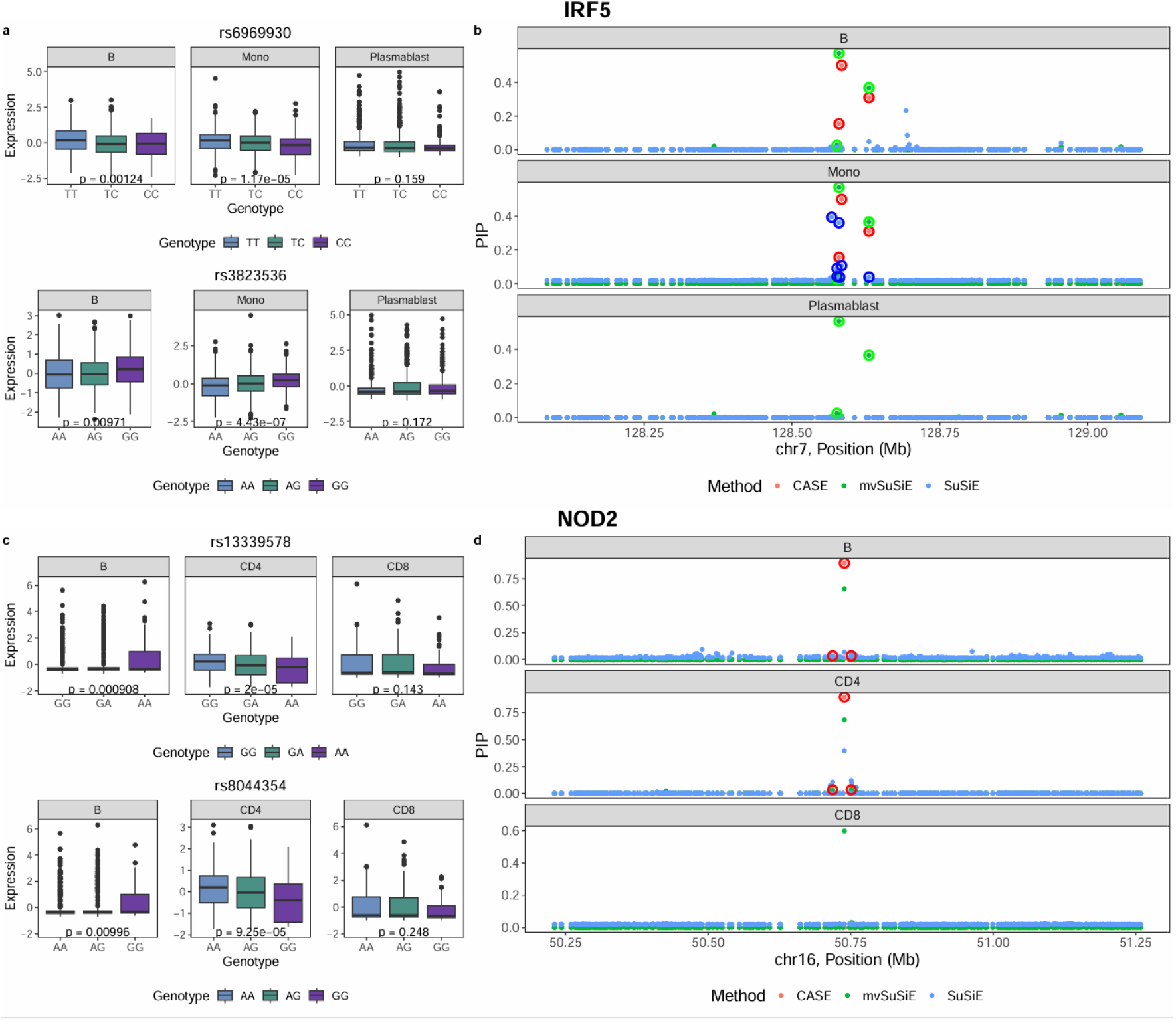
Examples of Fine-mapping results. **a-b**, Fine-mapping results in the OneK1K eQTL data for two genes, IRF5 (**a**) and NOD2 (**b**), using CASE (red), mvSuSiE (green), and SuSiE (blue). For each plot, we only displayed three typical cell types as examples. The left side shows boxplots exhibiting allelic directions on the adjusted expressions along with the nominal p-value for two lead variants in the credible sets. The right side shows the posterior inclusion probabilities (PIPs) for each variant among the three methods. The variants in credible sets are surrounded with circles.

The second gene, NOD2, is important in Crohn’s disease pathogenesis^35,36^. Among the three methods, only CASE identifies a three-SNP credible set in the B and CD4 cells, with lead SNPs rs13339578 and rs8044354 (Figs. 5c-d). These SNPs act as super-enhancers, and were supported by the significant marginal p-values in the corresponding cell types. Notably, these SNPs are validated in GTEx eQTL results of whole blood tissue with negative effect sizes. However, single-cell analysis exhibits opposite cell-type-specific effects in B and CD4 cells (Fig. 5c). In contrast, mvSuSiE and SuSiE lack the power to identify credible sets in any cell types. These results suggest that CASE outperforms mvSuSiE and SuSiE in signal detection for this gene by effectively leveraging cell type similarities.

### CASE identifies functionally enriched and disease-associated eGenes and eQTLs

To understand the biological functions and processes associated with eGenes identified by CASE, we performed enrichment analysis on hallmark pathways and three housekeeping gene lists derived from microarray data, bulk RNA-seq data, and single-cell RNA-seq data^37-39^. The analysis focused on three categories of eGenes: (1) *all-eGenes*: eGenes with at least one eQTL in at least one cell type), (2) *shared-eGenes*: eGenes with at least one eQTL in all cell types, and (3) *specific-eGenes*: eGenes with at least one eQTL in only a single cell type. For each category, we calculated the odds ratio relative to each gene list and applied Fisher’s exact test to assess statistical significance with BH p-value adjustment. We find that shared-eGenes are highly enriched across all three housekeeping gene lists, as well as in some cellular maintenance and metabolic pathways (Fig. 6a). These findings suggest that shared-eGenes are predominantly associated with fundamental cellular processes crucial for the survival and functionality of most cell types, highlighting their role in preserving essential biological activities that are universally required in various cellular environments.

**Fig. 6.**
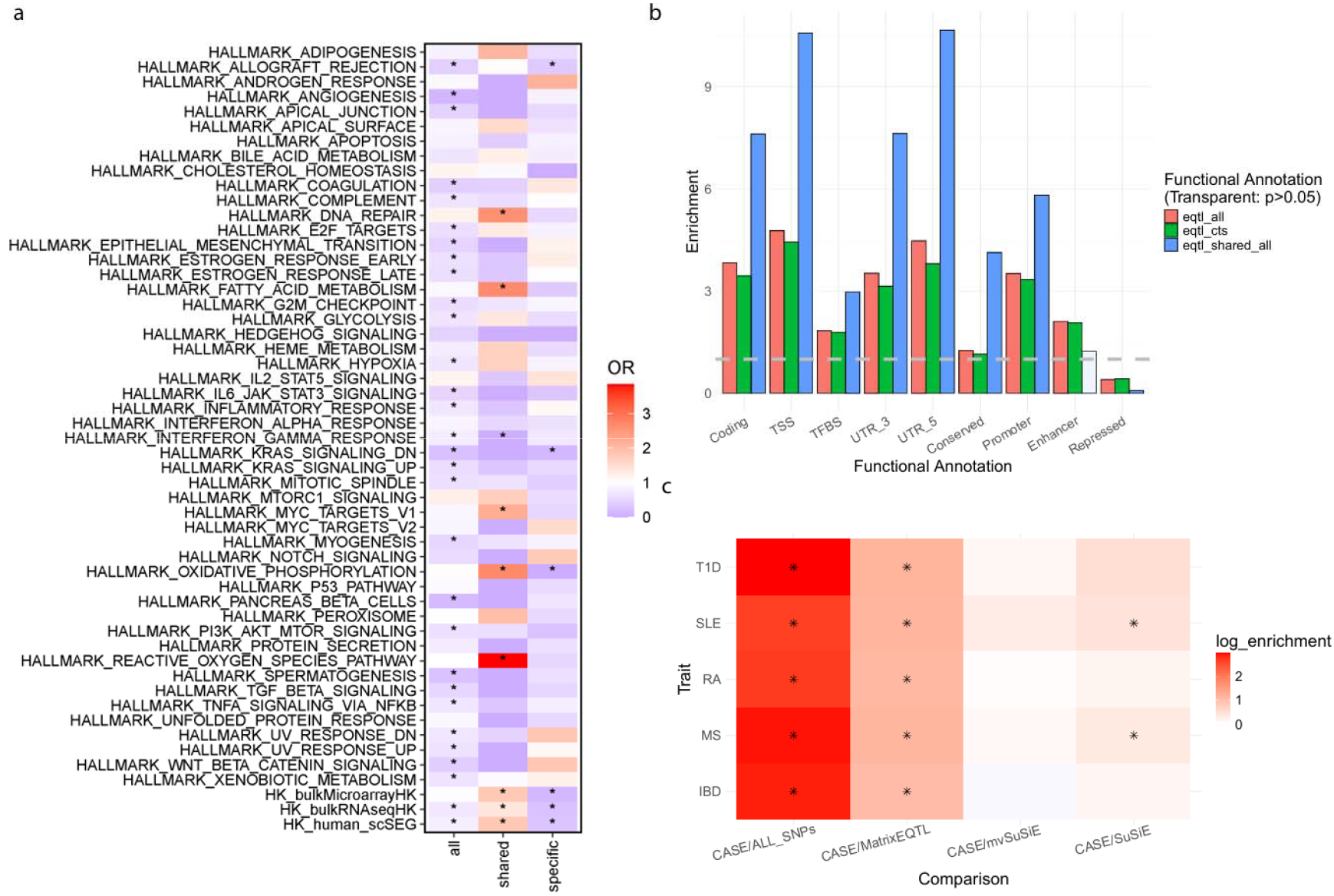
Enrichment analyses for eGenes and eQTLs. **a**, Heatmap of enrichment for different categories of eGenes: (1) *all-eGenes*, which have at least one eQTL in at least one cell type; (2) *shared-eGenes*, which have at least one eQTL in all cell types; and (3) *specific-eGenes*, which have at least one eQTL in only one cell type, against the hallmark pathways and three housekeeping gene lists. Colors represent odds ratio values, and asterisks indicate significance based on Fisher’s exact test with an adjusted p-value threshold of less than 0.2. **b**, Functional enrichment analysis for nine functional categories of SNPs. TSS: transcription start site; TFBS: transcription factor binding site. The SNPs are grouped by the effect-sharing categories of eQTLs: (1) *eQTL_all*: present in at least one cell type (red); (2) *eQTL_cts*: unique to one cell type (green); and (3) *eQTL_shared_all*: shared across all cell types (blue). Statistically nonsignificant (adjusted p-values > 0.05) results are shown in transparency. The grey line represents no enrichment (*y* = 1). **c**, Heritability enrichment analysis for auto-immune diseases. The y-axis represents diseases (*IBD*: Inflammatory Bowel Disease; *MS*: Multiple Sclerosis; *RA*: Rheumatoid Arthritis; *SLE*: Systemic Lupus Erythematosus; *T1D*: Type 1 Diabetes) and the x-axis represents comparisons between eQTLs identified by CASE and all SNPs, eQTLs identified by MatrixEQTL, eQTLs identified by mvSuSiE and eQTLs identified by SuSiE. We use asterisks for significant disease-category pairs (adjusted p-values <0.1).

To characterize the functional roles of eQTLs identified by CASE, we investigated functional enrichment across nine categories, including coding regions, TSS, transcription factor binding site (TFBS), 3′ UTR, 5′ UTR, conserved regions, promoter, enhancer and repressed regions. We examined three types of eQTLs: (1) present in any cell type, (2) shared among all cell types, and (3) unique to a single cell type. We applied Fisher’s exact test to assess statistical significance, with BH-adjusted p-values. Overall, eQTLs are generally depleted in the repressed regions but enriched in eight other active functional categories (Fig. 6b). Universally shared eQTLs demonstrate higher enrichment in functional categories associated with transcription, translation, and structural stability of genes, critical for core regulatory processes necessary for gene expression in all cell types. In contrast, cell-type-specific eQTLs are particularly enriched in enhancers, consistent with previous findings that highlight their role in modulating cell-type-specific gene expression by influencing the unique functional profiles of individual cell types^40^.

To explore the regulatory roles of eQTLs in complex diseases, we utilized GWAS summary statistics for five autoimmune diseases likely related to PBMCs, primarily from European population cohorts (Table 1). We compared the disease heritability enrichments of eQTLs identified by CASE, mvSuSiE, SuSiE, and MatrixEQTL (a marginal association test) using stratified LD score regression (S-LDSC)^41^ (Method). For all five diseases, CASE-identified eQTLs are significantly enriched compared to the baseline set of all SNPs and MatrixEQTL-identified ones (Fig. 6c), indicating the potential of eQTL fine-mapping analysis in prioritizing disease-related variants regulating gene expressions. While CASE-identified eQTLs exhibits higher heritability enrichments compared to mvSuSiE and SuSiE, the large overlap of eQTLs across the three methods rendered the differences statistically nonsignificant. We also investigated heritability enrichment across CASE-identified eQTLs with different sharing patterns. However, due to the limited number of eQTLs within each sharing category, the statistical power to detect significant heritability enrichment was limited. Nevertheless, the eQTLs shared among all cell types generally exhibit the highest enrichment (Supplementary Fig. 3), reflecting their likely involvement in core regulatory mechanisms underlying autoimmune diseases.

**Table 1.**
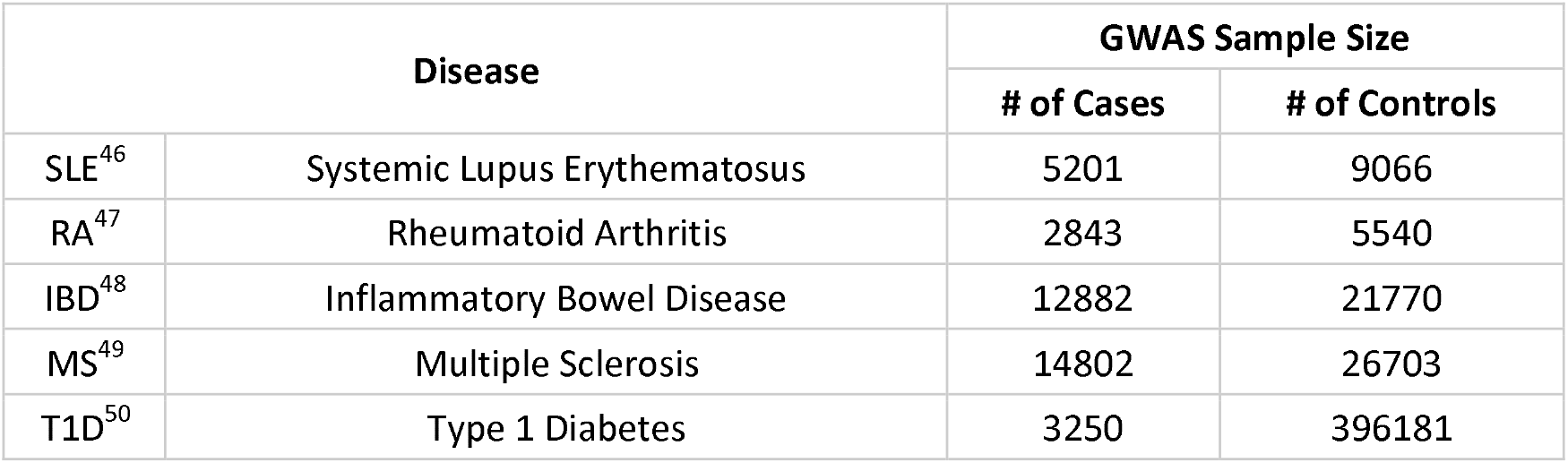
GWAS Summary Statistics of Autoimmune Diseases.

## Discussion

In this paper, we have proposed CASE, a novel, powerful and flexible framework for single-cell eQTL fine-mapping that enhances statistical power by effectively capturing cell type specificity and similarity. Compared to existing methods, CASE provides a flexible model capable of accommodating sharing patterns, minimizing overemphasis on fully shared effects, and identifying varied effect-sharing landscapes across cell types in the local genetic regions near genes. By appropriate characterizing cell-type specificity and similarity, CASE improves the statistical power in single-cell eQTL fine-mapping. Although tailored for single-cell eQTL fine-mapping, CASE is adaptable for broader applications, including multi-trait GWAS, multi-tissue eQTL, and multi-cell-type methylation QTL (mQTL) fine-mapping.

The application of CASE to the OneK1K dataset offers insights into complex disease mechanisms through shared genetic regulations on gene expression levels across cell types. CASE reveals novel eGenes and eQTLs at the cell type level, highlighting the importance of analyzing single-cell data. Unlike the original OneK1K analysis, where most identified eQTLs are unique to individual cell type, CASE identifies a significantly higher number of shared effects across cell types. Comparisons between cell-type-specific and shared eQTLs and eGenes in terms of functionality and pathways suggest the different roles of cell-type-specific and shared eQTLs and eGenes. In brief, universally shared eQTLs play a crucial role in fundamental cellular functions across all cell types, while cell-type-specific eQTLs uniquely contribute to the specialized functional profiles of individual cell types.

Despite the many improvements of CASE over existing methods, several limitations remain. First, CASE results still reflect a common challenge in eQTL studies that the number of eGenes and eQTLs are highly correlated with sample size and cell type proportions, as observed in prior studies^13,42^. As a result, achieving sufficient statistical power in rare cell types remains challenging with limited data. To study those rare cell types, a larger sample size is expected. Second, the current implementation of CASE relies on the accuracy of the LD estimates. Therefore, if the external LD panel differs from the study population, potentially due to ancestry differences, allele flipping or other issues, CASE may infer incorrect eQTLs. Incorporating LD calibration tools like CARMA^43^ could help address these inconsistency issues, improving CASE’s robustness against LD mismatches. Third, CASE estimates the effect-sharing patterns solely from the data without any prior knowledge. However, recent studies suggest that integrating functional annotations to the priors may improve fine-mapping methods^21,43,44^, which could prioritize causal eQTLs to fit the priors and enhance accuracy and efficiency in identifying causative regulatory variants. Lastly, although CASE has a comparable runtime to mvSuSiE, it still needs numerous MCMC iterations for (Supplementary Fig. 4). Considering that CASE primarily improves other existing methods through the prior fitting stage, we could update the posterior calculation step using a faster algorithm like variational inference^45^, an approach widely used in other fine-mapping methods, to alleviate computation demands.

## Acknowledgements

This work was supported in part by the National Institutes of Health R01 GM134005, U24 HG012108, and U01 HG013840. We thank Yazar *et al*. for sharing the OneK1K data.

## Data availability

OneK1K single-cell gene expression and genotype data are available via Gene Expression Omnibus (GSE196830). The cell-by-gene data are available at Human Cell Atlas (HCA) (https://cellxgene.cziscience.com/e/3faad104-2ab8-4434-816d-474d8d2641db.cxg). The GTEx v10 eQTL summary statistics are available at the GTEx Portal (https://gtexportal.org/home/downloads/adult-gtex/qtl). The availability of auto-immune disease GWAS summary statistics are summarized in Table 1. The functional annotations of SNPs are from the BaselineLD v.2.1 (https://data.broadinstitute.org/alkesgroup/LDSCORE/).

## Code availability

Software implementing CASE can be found at https://github.com/leaffur/CASE.

## Author Contributions

C.L. and H.Z. conceived the study and developed the statistical model. C.L. performed statistical analysis. W.L. assisted in eQTL analysis for OneK1K data. X.Z. assisted in fine-mapping analysis for OneK1K data. L.X. prepared GWAS summary statistics for auto-immune diseases. C.L. and Y.L. performed enrichment analyses. H.Z. advised on statistical and genetic analysis. C.L. implemented the software. All authors contributed to manuscript writing and editing and approved the manuscript.

## Methods

### Model and Assumptions

For a given gene, we consider the genetic effects of SNPs between 500Kb upstream and downstream of the transcription start site (TSS) as its *cis*-region, and we use *M* to denote the number of *cis*-SNPs. Suppose there are *C* cell types, each with sample size *N*_*C*_. For a given gene with *M* SNPs in its *cis*-region, the genetic regulation of covariate-adjusted expression levels across cell types can be formulated as the following multivariate multiple regression model:

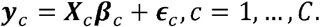

Here, 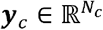 denotes the standardized gene expressions in cell type *c* after appropriate normalization and covariate adjustments (e.g., sex, age, genotype principal components (PCs), and gene expression PCs), 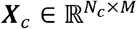 denotes the standardized genotypes within the *cis*-region of the gene, ***β***_*C*_ ∈ ℝ^*M*^ represents the *cis*-eQTL effect size, and 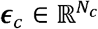 is the noise.

In single-cell eQTL studies, each pair of two cell types may share a majority of individuals. Therefore, we assume that the noise terms are independent among individuals for a certain cell type but correlated for the overlapped individuals across cell types. The noise is distributed from a multivariate normal distribution with mean zero and covariances:

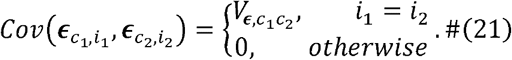

Overall, the covariance matrix of noise, ***V*** _*ϵ*_ ∈ ℝ^*C*× *C*^, in shared individuals across cell types can be expressed as

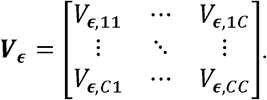

We denote the matrix of *cis*-eQTL effect sizes as ***B*** = (***β***_1_,…, ***β***_*C*_) ∈ ℝ^*M*×*C*^, and assume that each row ***B***_*j*_·, representing the effect size of the SNP j across cell types, follows a mixture normal distribution with T components:

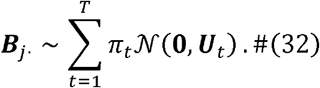

Here, 𝒩 (·) denotes a multivariate normal distribution, each covariance matrix ***U***_*t*_ ∈ ℝ^*C*× *C*^ characterizes a specific pattern of shared effect sizes, and π_*t*_ indicates the probability of each pattern. For example, if no effect exists in cell type c under a pattern ***U***_*t*_, the cth row and column of ***U***_*t*_ contain only zeros. The choice of the number of patterns, *T*, will be discussed later. The last pattern ***U***_*T*_ is fixed zero, representing SNPs with no eQTL effect in any cell type. The covariance of ***B***_*j*_· is 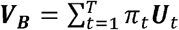. We further assume independence among the *cis*-eQTL effect size, noise and genotype. Then the phenotypic covariance matrix of the gene expressions can be expressed as ***V***_*y*_ = ***MV***_***B***_ + ***Vϵ*** (Supplementary Notes).

Let 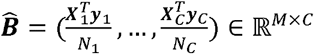 denote the eQTL summary statistics from gene-SNP marginal association tests. Given the true *cis*-eQTL effect size, 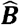 asymptotically follows a matrix normal distribution (Supplementary Notes):

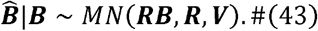

Here, *MN*(·) represents a matrix normal distribution, ***V*** is the sample-adjusted covariance of gene expression with 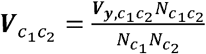, where 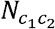 denotes the overlapping sample size between cell type *C*_1_ and *C*^2^, and ***R*** ∈ ℝ^*M*×*M*^ represents the LD matrix, which can be estimated from the study population or reference panels.

To learn the true eQTL effect sizes based on the observed summary statistics, we apply an Empirical Bayesian framework. The prior is given from our assumptions in Equations (1) and (2), where the parameters in the prior can be estimated from the observed summary statistics, and the distribution of the data given the fitted prior is provided in Equation (3). An overview of the CASE approach is:

1. Estimate the sample-adjusted total covariance, 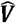, using weak signals from eQTL summary statistics.
2. Estimate the effect-sharing patterns and probabilities, 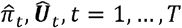, using a Monte-Carlo Expectation-Maximization (MCEM) algorithm.
3. Compute the posterior inclusion probability, 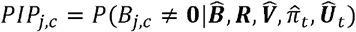, for each SNP j in cell type c, using a Markov chain Monte Carlo (MCMC) algorithm.
4. Obtain the credible sets (defined below) based on the posterior probabilities.

We discuss each step in detail in the following.

### Estimate the sample-adjusted covariance

There are several choices of the phenotypic covariance ***V***. A simple approach is to ignore the correlations and fix the off-diagonal elements at zero, which works for cell types with low phenotypic correlations. To quantify the phenotypic correlations, one approach is to apply existing methods like LD score regression (LDSC)^51^ among the overlapping samples for each cell-type pair. However, the computation time is heavy when the cell type number is large. In the CASE framework, we adopt a moment-based approach like mashr^18^, using the fact that for any matrix normal distribution, *X* ∼ *MN*(***µ, U, V***), 𝔼 (*X*− µ)^*T*^(*X*− µ) = ***V****Tr*(***U***). Let J{denote a set of marginally insignificant SNPs. We can assume that each SNP in ℋ is neither an eQTL nor in high LD with any eQTL, i.e., 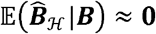. Thus, we have 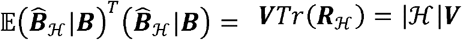, where 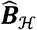 and ***R***_ℋ_ are summary statistics and LD matrix within the subset ℋ, and ℋ denotes the number of SNPs in the subset ℋ. We can estimate the sample covariance, 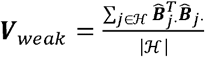, and scale it to obtain 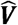.

### Estimate the effect-sharing patterns and probabilities using an MCEM algorithm

We are supposed to optimize the likelihood, 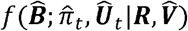, to fit the prior estimates, Θ= {π_*t*_,***U***_*t*_, *t* = 1,…, *T*}. Our proposal is to apply an EM algorithm^52^ for the optimization. A simpler version of this problem can be solved through an extreme deconvolution algorithm^30^, where each observation (SNP) is treated independently, i.e., ***R***= ***I***_*M*_. However, incorporating the LD adds more complexities to the algorithm.

Here we treat the true effects, ***B***_*j*·_, and its pattern or sharing, *g*_*j*_, as “missing variables”, where *g*_*j*_ =t ⇔ ***B***_*j·*_ ∼ 𝒩 (**0, *U***_*t*_). Then, the complete log likelihood for (***B, g***) after removing constant terms is:

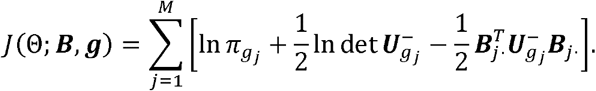

Here, 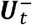 denotes the Moore–Penrose inverse of a matrix, as U may not be of full rank. In the *r*th iteration, the E-step takes the expectation of the complete log likelihood given the observed 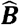 and current model parameters Θ^r^, which leads to

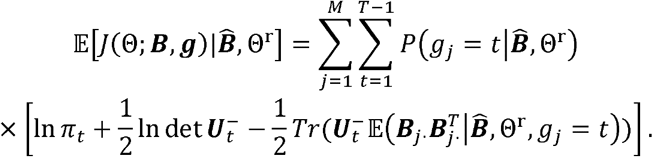

In the M-step, we obtain the formula to iteratively update the parameters which monotonically maximize the objective function (Supplementary Notes):

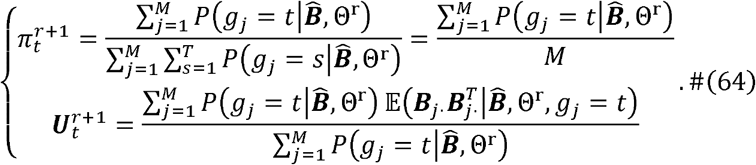

The conditional probabilities and expectations in Equation (4) can be quite complicated, because SNPs in the same LD block exhibit dependencies in the effect-sharing patterns. Considering the heavy complexity, we propose to use an MCMC algorithm to sample from the conditional distributions instead of using an explicit form. We adopt an exponential growth of the MC sample size following the suggestions from classical MCEM algorithms to ensure convergence^27-29^.

For computation concern, we only include SNPs with marginal FDR-adjusted p-values <= 0.2 in at least one cell type during the fitting process, with the assumption that marginally insignificant SNPs cannot be eQTLs. The reason is that the non-eQTL covariance matrix is fixed (***U***_*T*_ = **0**), and there is no need to include excessive data to fit a zero matrix. Note that we do not threshold SNPs in the following testing step, as our interests in the testing step transfer from learning the overall sharing pattern to identifying the eQTL effect size for each SNP. Any sharing pattern with π *=0* will be removed from the EM iteration, meaning that the number of sharing patterns *T* is implicitly and adaptively fitted.

### The initialization of the sharing patterns, *T*, π_*t*_ and *Ut*, in the MCEM algorithm

In practice, any kind of sharing pattern can be chosen for the initialization of *T*, π _*t*_ and ***U***_t_, while we make some recommendations to initialize the parameters based on the data. For a type of sharing pattern *t*, we call the cell types, {*c*_1_, *c*_2_,*…, c*_K_}, which have shared eQTL effects as “sharing cell types”, and the remaining cell types without eQTL effects as “non-sharing cell types”. We first introduce our approach to initialize ***u***_t_ based on the eQTL summary statistics, 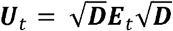, where *E*_t_ is a correlation matrix with ones on the diagonal and *ρ* on the off-diagonal for “sharing cell types”, and zeroes for “non-sharing cell types”, and ***D*** is a diagonal matrix with the cell-type-specific variances. Note that the parameter update Equation (4) of the MCEM algorithm preserves the rank and zero elements for each ***u***_t_, so the explicit zeroes correlations in ***E***_t_ for the non-sharing cell types is critical. By default, we fix *ρ = 0*.*1* and use the maximum of effects for 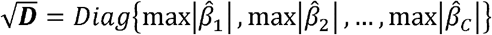, while the algorithm will iteratively estimate the magnitude of variances and correlations for the sharing cell types.

Next, we introduce our choice to initialize the mixture probability, π _*t*_. We calculate the probability of observing a sharing pattern *t* determined by the marginal testing p-values (after appropriate multiple testing adjustment). For example, if the sharing cell types are {*c*_1_, *c*_2_,…, *c*_*K*_}, we initialize π_*t*_ = *P*(*p*_1_ ≤ α,*p*_*r*_ > α,; *l* ∈ {*c*_l_, *c*_2_,…, *c*_*K*_},∉ {*c*_l_, *c*_2_,…, *c*_K_}), where in our implementation, *p* is the FDR-adjusted p-value for each variant, and α= 0.1 is the significance level.

Since the initialization approach of π_*t*_ is based on the observed significance, some sharing patterns may have π_*t*_ = 0, meaning that the given eQTL summary statistics do not support the existence of those sharing patterns. The non-zero π_*t*_’s implicitly determines the total number of sharing pattern, *T*. Considering the observed significance may be sensitive to the choice of p-value adjustment and significance level, a positive value can be added to some π_*t*_’s where we would like not to miss out on those sharing patterns during the initialization step. In our implementation, we add 1/*M* to π_*t*_ ‘s corresponding to each cell-type-specific pattern and a shared-by-all pattern, and re-normalize π_*t*_’s to sum to one.

### Sample true effects given the prior estimates using an MCMC algorithm

We propose coordinate-wise updates to iteratively sample from 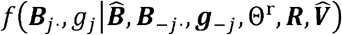 per SNP, reducing the model complexity from a joint sampling. At each iteration *i*, we loop for each SNP to sample the true effects. For the Jth SNP, we first sample a type of sharing pattern for 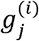 from

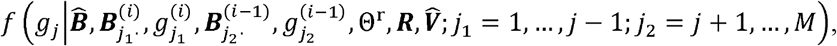

and then sample the effect from a normal distribution specific to the sharing pattern 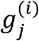,

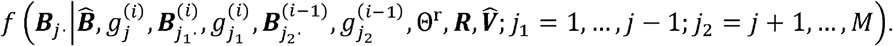

The details of the sampling distributions and algorithm can be found in the Supplementary Notes. Considering the computational burden, we thresholds the SNPs with extremely low PIPs or effect sizes to be zero in each iteration. We repeat the iteration until having enough MC samples. The MC samples (after removing burn-in samples in the first several iterations) are used to infer the PIPs and expectations of the true effects. This approach is applied for both prior fitting and posterior analysis steps.

The MCMC coordinate-wise updates may be sensitive to the initialization of B, g. To avoid this issue, we run MCMC multiple times when calculating the posteriors, with different random initialization each time^31,32^. The prior fitting step may not be affected by the sensitivity issue, as we are not interested in inferring the distribution of any specific SNP, but using the MC samples to fit the overall sharing patterns.

### Obtain the credible sets

Once we obtain the PIPs, a direct report of causal eQTLs is to use the SNPs with *PIP*_*j,c* ≥_1 − α, where α is some threshold of the significance level (e.g., α = 0.05). However, when one or more potential eQTLs occur in a high LD region, it is hard to distinguish which SNP(s) are true eQTL(s). In an extreme case, if a SNP is in perfect LD with an eQTL, we will ideally obtain *PIP*_*j,c*_ = 0.5 for both SNPs. To improve the LD-raised power issue, we adopt the idea to aggregate SNPs into credible sets cell type by cell type^5^. A credible set is the smallest subset of correlated variants, 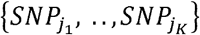, containing at least one causal variant which satisfies: 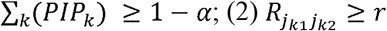, with default values α = 0.05 and *r*= 0.5.

### Simulations

Gene expression data were simulated using the genotyped OneK1K cohort (See Data Availability) across five distinct cell types with complex sharing patterns. For each gene, 1,000 contiguous SNPs were randomly selected from the *cis*-region on chromosome 2, from which 1 to 5 SNPs were chosen as eQTLs with probabilities 0.4, 0.25, 0.15, 0.1, and 0.1 respectively, reflecting *cis*-eQTL sparsity^1,42^. Each eQTL was assigned a random sharing pattern and sampled from a multivariate normal distribution with mean zero and a covariance matrix ***U***_*t*_. Finally, all eQTL effect sizes were scaled so that the *cis*-heritability for the five cell types is 0.1, 0.08, 0.07, 0.05, and 0.04, respectively. The heritability multiplier was fixed at one when varying the number of individuals in {400, 600, 800, 1000}, and conversely, the sample size was fixed at 1,000 when varying the heritability multiplier in {0.75, 1, 1.25, 1.5}.

The first simulated scenario reflected real genomic complexity. Each true eQTL was randomly selected to be cell-type-specific or shared among a random number of cell types with equal probability, with effect size correlations of 0.3 among shared cell types. We ran 100 simulations for each scenario and simulated 250 genes in each run. The second scenario was designed to investigate false positives arising from over-sharing. Two specific cases were considered: (1) over-sharing due to shared-by-all patterns, with cis-heritability values set at {0.1, 0.08, 0.07, 0.05, 0}; and (2) over-sharing due to shared-by-some patterns, with heritability values set at {0.1, 0.08, 0, 0, 0}. Both cases used a fixed sample size of 1,000 and simulated 200 genes. False positive rates were calculated as the fraction of falsely identified eGenes over the total number of non-eGenes (heritability equal to zero) across cell types. The significance level was varied from 0 to 0.5, corresponding to the coverage threshold varying from 1 to 0.5 when aggregating SNPs into credible sets. Since mvSuSiE does not provide cell-type-wise credible sets, we calculated the credible sets for mvSuSiE based on their local false discovery rate (lfsr).

### eQTL analysis in OneK1K

For the OneK1K dataset, we consolidated the original 29 cell sub-types into eight primary cell types based on cell differentiation patterns and clustering in Yazar *et al*.^13^, and removed cell types with fewer than 2,000 cells, as summarized in Supplementary Table 1. The single-cell Unique Molecular Identifier (UMI) counts for each cell type were aggregated per gene per individual to generate pseudo-bulk expression profiles. Genes expressed in fewer than 10% of individuals were excluded from eQTL analysis. For each cell type, pseudo-bulk counts were normalized using the trimmed mean of M-values method (TMM)^53^ and log2-transformed counts per million mapped reads (CPM) were computed using edgeR^54^. We further removed genes with TSS in the extended MHC region (2.5-3.4 Mb on chromosome 6), due to the long-range LD and potential extreme values in summary statistics in the region^55,56^. Biallelic, autosomal SNPs were filtered to include SNPs with a minor allele frequency (MAF) greater than 5%, Hardy-Weinberg equilibrium *P*>1 × 10^−6^, and further pruned to remove highly correlated SNPs (−-indep-pairwise 250 50 0.95) using plink2^57^.

The *cis*-eQTL summary statistics were obtained using MatrixEQTL^58^ for each cell type separately, adjusting for potential confounders including age, sex, the first two genotype PCs, and the first two expression PCs. Only SNPs between 500Kb upstream and downstream of each gene’s TSS were included in the eQTL analysis. For genes unexpressed in certain cell types (or removed due to extremely low expressions as previously mentioned), missing z-scores were replaced with zero in the summary statistics. The reference LD was derived from the OneK1K in-sample genotypes. For statistical fine-mapping analysis, we applied CASE, mvSuSiE, and SuSiE only for genes with at least one significantly associated SNP (FDR ≤ 0.2) in at least one cell type, yielding 11,704 genes to be analyzed. To improve interpretability, we excluded SNPs with extremely low posterior inclusion probability (PIP ≤ 0.05) when analyzing the eQTLs.

### Quality control on the GWAS summary statistics for auto-immune diseases

To ensure robust analysis, we conducted quality control on the GWAS summary statistics for each trait following LDHub guidelines^59^. Specifically, we removed SNPs with effective sample sizes less than 0.67 times the 90th percentile of the sample size, as well as insertions and deletions (INDELs), structural variants, and strand-ambiguous (A/T and G/C) SNPs using the LD score regression (LDSC) software^60^.

### Gene marker enrichment analysis across cell types

We performed an enrichment analysis of cell-type-specific eGenes against a list of marker genes derived from the OneK1K scRNA-seq data, generated using COSG^61^ with a fixed number of 200 marker genes per cell type. Given the high similarity among *T* subtypes and between DCs and monocytes, we aggregated *T* subtypes (CD4, CD8, and others) into a single *T*-cell category and combined DCs and monocytes into a myeloid category. For each cell type, we calculated the odds ratio between the marker genes and the cell-type-specific eGenes and assessed statistical significance using Fisher’s exact test with Benjamini-Hochberg (BH) p-value adjustment.

### Heritability enrichment analysis of auto-immune diseases

We used stratified LDSC (S-LDSC)^41^ to perform heritability enrichment analysis of auto-immune diseases. The LD scores were calculated with OneK1K genotypes using a 300-Kb LD window. For MatrixEQTL-derived eQTLs, we focused on SNPs with Bonferroni-adjusted *P* < 0.05. To study the enrichment of different sharing patterns, we used binary annotations categorizing eQTLs into overlapping groups: (1) present in any cell type, (2) shared among all cell types, (3) unique to a single cell type, (4) specific to T cells (shared or cell-type-specific among CD4, CD8 and other T cells), (5) specific to myeloid cells (shared or cell-type-specific among DC and Mono), and (6-13) specific to each individual cell type.

