## Supplementary for "Leveraging cell-type specificity and similarity improves single-cell eQTL fine-mapping"

### Supplementary Notes

#### Transform z scores to the estimated effects of standardized genotype and phenotype, $\hat{\boldsymbol{B}}$

In CASE model, we only discuss the situations where $\hat{\boldsymbol{B}}$ is estimated from the standardized genotypes and phenotypes. However, the accessible eQTL summary statistics may be z scores or estimates from unstandardized genotypes and phenotypes. Here we show the formula to transform the z scores to the required $\hat{\boldsymbol{B}}$.

We consider a simple case for a single SNP and single cell type. The matrix of summary statistics can be transformed element-wise. Suppose $\boldsymbol{x}, \boldsymbol{y} \in \mathbb{R}^N$ are standardized vectors, where $\boldsymbol{x}^T\boldsymbol{x} = \boldsymbol{y}^T\boldsymbol{y} = N - 1$. Regressing $\boldsymbol{y}$ on $\boldsymbol{x}$ results in

$$\hat{\beta} = \frac{\boldsymbol{x}^T\boldsymbol{y}}{\boldsymbol{x}^T\boldsymbol{x}} = \frac{\boldsymbol{x}^T\boldsymbol{y}}{N-1},$$

$$\widehat{var}(\hat{\beta}) = \frac{\left\|\boldsymbol{y} - \boldsymbol{x}\hat{\beta}\right\|^2}{N-1}(\boldsymbol{x}^T\boldsymbol{x})^{-1} = \frac{1-\hat{\beta}^2}{N-1}.$$

The standardization of predictors and responses does not affect the z scores for simple linear regressions, which means

$$z = \frac{\hat{\beta}}{\widehat{s.e.}(\hat{\beta})} = \hat{\beta}\sqrt{\frac{N-1}{1-\hat{\beta}^2}}.$$

Thus, we can approximately obtain $\hat{\boldsymbol{B}}$ by transforming the z scores element-wise:

$$\widehat{var}(\hat{\beta}) = \frac{1}{z^2 + N - 1},$$

$$\hat{\beta} = \sqrt{\frac{z^2}{z^2 + N - 1}}.$$

**The phenotypic variance**

Suppose $\boldsymbol{y}, \boldsymbol{\epsilon} \in \mathbb{R}^C$ and $\boldsymbol{x} \in \mathbb{R}^M$ are gene expressions, noise and genotype at the individual level, $\boldsymbol{B} \in \mathbb{R}^{M\times C}$ is the eQTL effect size matrix, and $\boldsymbol{R} \in \mathbb{R}^{M\times M}$represents the LD matrix. We have $\boldsymbol{y} = \boldsymbol{x}^T\boldsymbol{B} + \boldsymbol{\epsilon}$. The genotype, error and effect sizes are assumed to be mutually independent. Then, the phenotypic covariance matrix of the gene expressions can be expressed as

$$\begin{aligned}\boldsymbol{V_y} &= Cov(\boldsymbol{y}) \\ &= Cov(\boldsymbol{x}^T\boldsymbol{B} + \boldsymbol{\epsilon}) \\ &= Cov(\boldsymbol{x}^T\boldsymbol{B}) + Cov(\boldsymbol{\epsilon}) \\ &= \mathbb{E}(\boldsymbol{B}^T\boldsymbol{x}\boldsymbol{x}^T\boldsymbol{B}) + \boldsymbol{V_\epsilon} \\ &= \mathbb{E}(\boldsymbol{B}^T\boldsymbol{R}\boldsymbol{B}) + \boldsymbol{V_\epsilon} \\ &= M\boldsymbol{V_B} + \boldsymbol{V_\epsilon}.\end{aligned}$$

The last equality holds as for each pair of cell types, $c_1, c_2$, $\mathbb{E}\left(\boldsymbol{\beta}_{c_1}^T\boldsymbol{R}\boldsymbol{\beta}_{c_2}\right) = Tr\left(\mathbb{E}\left(\boldsymbol{\beta}_{c_2}\boldsymbol{\beta}_{c_1}^T\right)\cdot\boldsymbol{R}\right) = Tr\left(V_{B,c_1c_2}\boldsymbol{I}_M\cdot\boldsymbol{R}\right) = MV_{\boldsymbol{B},c_1c_2}$.

**Derive the conditional distribution of eQTL summary statistics given true effects**

In this section, we show a brief proof for $\widehat{\boldsymbol{B}}|\boldsymbol{B} \sim MN(\boldsymbol{RB}, \boldsymbol{R}, \boldsymbol{V})$**.** The single-trait form ($C = 1$) has been well studied in previous research[1-3] that $\widehat{\boldsymbol{\beta}}|\boldsymbol{\beta} \to_d \mathcal{N}(\mathrm{R}\boldsymbol{\beta}, \boldsymbol{R}/N)$. Let $vec(\cdot)$ denote the vectorization of a matrix (by column). The distribution of $vec(\widehat{\boldsymbol{B}})|\boldsymbol{B}$ can be inferred from the same format of the single-trait case that $vec\left(\widehat{\boldsymbol{B}}\right)|\boldsymbol{B}$ is asymptotically normal. The proof can be found in X. Zhu and M. Stephens[27], majorly applying the central limiting theorem and the delta method. The main difference for the multi-trait form is that the covariance term is more complicated due to the covariances across cell types. Here, we focus on the values of the mean and covariance in the asymptotic distribution.

We first obtain the expectation of each column of $\widehat{\boldsymbol{B}}$, $\widehat{\boldsymbol{\beta}}_c$:

$$\mathbb{E}(\widehat{\boldsymbol{\beta}}_c|\boldsymbol{\beta}_c) = \mathbb{E}\left(\frac{\boldsymbol{X}_c^T\boldsymbol{y}_c}{N_c}|\boldsymbol{\beta}_c\right)$$

$$= \mathbb{E}\left(\frac{\boldsymbol{X}_c^T\boldsymbol{X}_c\boldsymbol{\beta}_c}{N_c}|\boldsymbol{\beta}_c\right) + \mathbb{E}\left(\frac{\boldsymbol{X}_c^T\boldsymbol{\epsilon}_c}{N_c}|\boldsymbol{\beta}_c\right)$$

$$= \boldsymbol{R\beta}_c.$$

Thus, $\mathbb{E}(\widehat{\boldsymbol{B}}|\boldsymbol{B}) = \boldsymbol{RB}$, and $\boldsymbol{\mu} = \mathbb{E}(vec(\widehat{\boldsymbol{B}})|\boldsymbol{B}) = vec(\boldsymbol{RB})$.

Next, we obtain the covariances for each pair of columns in $\widehat{\boldsymbol{B}}$. For simplicity, we assume that only the first $N_{c_1c_2}$ individuals are shared for cell type $c_1$ and $c_2$. (Here we allow the case that $c_1 = c_2$, where $N_{c_1c_2} = N_{c_1}$). We use a conclusion from Zhang, Y. *et al.*[4] that $\mathbb{E}(\boldsymbol{X}_{c_1}^T\boldsymbol{X}_{c_2}\boldsymbol{X}_{c_1}^T\boldsymbol{X}_{c_2}) \approx N_{c_1}N_{c_2}\boldsymbol{R}^2 + MN_{c_1c_2}\boldsymbol{R}$. Provided the sparsity of the eQTL, and $\mathbb{E}\left(\boldsymbol{\beta}_{c_1}\boldsymbol{\beta}_{c_2}^T\right) = V_{B,c_1c_2}\boldsymbol{I}$, we have $\boldsymbol{\beta}_{c_1}\boldsymbol{\beta}_{c_2}^T = V_{B,c_1c_2}\boldsymbol{I} + o(\boldsymbol{V_B})$.

$$cov(\widehat{\boldsymbol{\beta}}_{c_1}, \widehat{\boldsymbol{\beta}}_{c_2}|\boldsymbol{B}) = \mathbb{E}(\widehat{\boldsymbol{\beta}}_{c_1}\widehat{\boldsymbol{\beta}}_{c_2}^T|\boldsymbol{B}) - \mathbb{E}(\widehat{\boldsymbol{\beta}}_{c_1}|\boldsymbol{B})\mathbb{E}(\widehat{\boldsymbol{\beta}}_{c_2}|\boldsymbol{B})^T$$

$$= \mathbb{E}\left(\frac{\boldsymbol{X}_{c_1}^T\boldsymbol{X}_{c_1}\boldsymbol{\beta}_{c_1}\boldsymbol{\beta}_{c_2}^T\boldsymbol{X}_{c_2}^T\boldsymbol{X}_{c_2} + \boldsymbol{X}_{c_1}^T\boldsymbol{\epsilon}_{c_1}\boldsymbol{\epsilon}_{c_2}^T\boldsymbol{X}_{c_2}}{N_{c_1}N_{c_2}}|\boldsymbol{B}\right) - \boldsymbol{R\beta}_{c_1}\boldsymbol{\beta}_{c_2}^T\boldsymbol{R}$$

$$= \frac{1}{N_{c_1}N_{c_2}}\mathbb{E}(\boldsymbol{X}_{c_1}^T\boldsymbol{X}_{c_1}(V_{B,c_1c_2}\boldsymbol{I} + o(\boldsymbol{V_B}))\boldsymbol{X}_{c_2}^T\boldsymbol{X}_{c_2}|\boldsymbol{B}) + \frac{N_{c_1c_2}V_{\epsilon,c_1c_2}}{N_{c_1}N_{c_2}}\boldsymbol{R} - \boldsymbol{R}(V_{B,c_1c_2}\boldsymbol{I} + o(\boldsymbol{V_B}))\boldsymbol{R}$$

$$\approx \frac{V_{B,c_1c_2}}{N_{c_1}N_{c_2}}\mathbb{E}(\boldsymbol{X}_{c_1}^T\boldsymbol{X}_{c_1}\boldsymbol{X}_{c_2}^T\boldsymbol{X}_{c_2}) + \frac{N_{c_1c_2}V_{\epsilon,c_1c_2}}{N_{c_1}N_{c_2}}\boldsymbol{R} - V_{B,c_1c_2}\boldsymbol{R}^2$$

$$\approx \frac{N_{c_1c_2}MV_{B,c_1c_2}}{N_{c_1}N_{c_2}}\boldsymbol{R} + \frac{N_{c_1c_2}V_{\epsilon,c_1c_2}}{N_{c_1}N_{c_2}}\boldsymbol{R}$$

$$= \frac{N_{c_1c_2}V_{y,c_1c_2}}{N_{c_1}N_{c_2}}\boldsymbol{R}.$$

Thus, we have $\boldsymbol{Cov}\left(vec(\widehat{\boldsymbol{B}})\middle|\boldsymbol{B}\right) \approx \boldsymbol{V} \otimes \boldsymbol{R}$.

Therefore, we conclude that $vec(\widehat{\boldsymbol{B}})|\boldsymbol{B} \rightarrow_d \mathcal{N}(vec(\boldsymbol{RB}), \boldsymbol{V} \otimes \boldsymbol{R})$, i.e., $\widehat{\boldsymbol{B}}|\boldsymbol{B} \rightarrow_d MN(\boldsymbol{RB}, \boldsymbol{R}, \boldsymbol{V})$. The result is consistent with the single-trait form, because the diagonal elements are $1/N_c$ in $\boldsymbol{V}$ due to the standardized gene expressions.

**Optimizing the Expected likelihood in the EM algorithm**

In the M-step, we want to optimize the expected likelihood,

$$\mathbb{E}\left[J(\Theta; \boldsymbol{B}, \boldsymbol{g})|\widehat{\boldsymbol{B}}, \Theta^{\mathrm{r}}\right] = \sum_{j=1}^{M}\sum_{t=1}^{T-1} P\left(g_j = t\middle|\widehat{\boldsymbol{B}}, \Theta^{\mathrm{r}}\right)$$
$$\times \left[\ln \pi_t + \frac{1}{2}\ln\det \boldsymbol{U}_t^{-} - \frac{1}{2}Tr(\boldsymbol{U}_t^{-}\mathbb{E}\left(\boldsymbol{B}_{j\cdot}\boldsymbol{B}_{j\cdot}^{T}\middle|\widehat{\boldsymbol{B}}, \Theta^{\mathrm{r}}, g_j = t\right))\right].$$

Here, we introduce some necessary equations without proof: (1) $\pi_T = 1 - \sum_{t=1}^{T-1}\pi_t$ ; (2) $\partial \ln\det \boldsymbol{X} / \partial \boldsymbol{X} = (\boldsymbol{X}^{-1})^T$; (3) $\partial Tr(\boldsymbol{AB})/\partial \boldsymbol{A} = \boldsymbol{B}^T$. Then, we calculate the first derivatives of the objective function with respect to $\pi_t$ and $U_t^{-}$, $t = 1, \dots, T-1$:

$$\begin{cases} \dfrac{\partial J}{\partial \pi_t} = \dfrac{\sum_{j=1}^{M} P\left(g_j = t\middle|\widehat{\boldsymbol{B}}, \Theta^{\mathrm{r}}\right)}{\pi_t} - \dfrac{\sum_{j=1}^{M} P\left(g_j = T\middle|\widehat{\boldsymbol{B}}, \Theta^{\mathrm{r}}\right)}{\pi_T} = 0 \\ \dfrac{\partial J}{\partial \boldsymbol{U}_t^{-}} = \dfrac{1}{2}\boldsymbol{U}_t \sum_{j=1}^{M} P\left(g_j = t\middle|\widehat{\boldsymbol{B}}, \Theta^{\mathrm{r}}\right) - \dfrac{1}{2}\sum_{j=1}^{M} P\left(g_j = t\middle|\widehat{\boldsymbol{B}}, \Theta^{\mathrm{r}}\right)\mathbb{E}\left(\boldsymbol{B}_{j\cdot}\boldsymbol{B}_{j\cdot}^{T}\middle|\widehat{\boldsymbol{B}}, \Theta^{\mathrm{r}}, g_j = t\right) = \boldsymbol{0} \end{cases}.$$

Correspondingly, we obtain the formula to iteratively update the parameters which monotonically maximize the objective function:

$$\begin{cases} \pi_t = \dfrac{\sum_{j=1}^{M} P\left(g_j = t \middle| \widehat{\boldsymbol{B}}, \Theta^{\mathrm{r}}\right)}{\sum_{j=1}^{M} \sum_{s=1}^{T} P\left(g_j = s \middle| \widehat{\boldsymbol{B}}, \Theta^{\mathrm{r}}\right)} = \dfrac{\sum_{j=1}^{M} P\left(g_j = t \middle| \widehat{\boldsymbol{B}}, \Theta^{\mathrm{r}}\right)}{M} \\ \boldsymbol{U}_t = \dfrac{\sum_{j=1}^{M} P\left(g_j = t \middle| \widehat{\boldsymbol{B}}, \Theta^{\mathrm{r}}\right) \mathbb{E}\left(\boldsymbol{B}_{j\cdot} \boldsymbol{B}_{j\cdot}^T \middle| \widehat{\boldsymbol{B}}, \Theta^{\mathrm{r}}, g_j = t\right)}{\sum_{j=1}^{M} P\left(g_j = t \middle| \widehat{\boldsymbol{B}}, \Theta^{\mathrm{r}}\right)} \end{cases}.$$

<u>**Algorithm 1** MCEM: Fitting the prior**:**</u>

**Require:** eQTL summary statistics, $\widehat{\boldsymbol{B}}$, LD matrix $\boldsymbol{R}$, sample-adjusted covariance matrix $\widehat{\boldsymbol{V}}$.

Initialize the prior parameters, $\pi_t^{(0)}, \boldsymbol{U}_t^{(0)}, t = 1, \cdots, T$.

**Repeat**

MC-step: Generate $W$ samples of eQTL effect size matrices, $\boldsymbol{B}^{(w)}$, and the hidden indicators, $\boldsymbol{g}^{(w)}$, $w = 1, \cdots, W$, from the posterior distributions given the current $\pi_t^{(r)}, \boldsymbol{U}_t^{(r)}$ using the MCMC algorithm (See below).

E-step: Obtain the posterior expectations with the MC samples:

$$p_{kt} = P\left(g_j = t \middle| \widehat{\boldsymbol{B}}, \pi_t^{(r)}, \boldsymbol{U}_t^{(r)}\right) = \frac{\sum_w 1_{\left\{g_k^{(w)} = t\right\}}}{\sum_{w,t} 1_{\left\{g_k^{(w)} = t\right\}}},$$

$$\mathbf{S}_{kt} = \mathbb{E}\left(\boldsymbol{B}_j \boldsymbol{B}_j^T \middle| g_j = t, \widehat{\boldsymbol{B}}, \pi_t^{(r)}, \boldsymbol{U}_t^{(r)}\right) = \frac{\sum_w 1_{\left\{g_k^{(w)} = t\right\}} \boldsymbol{B}_j^{(w)} \boldsymbol{B}_j^{(w),T}}{\sum_{w,t} 1_{\left\{g_k^{(w)} = t\right\}}}.$$

M step: Update the prior parameter:

$$\pi_t^{(r+1)} = \frac{\sum_k p_{kt}}{\sum_{k,t} p_{kt}},$$

$$\boldsymbol{U}_t^{(r+1)} = \frac{\sum_k p_{kt} \mathbf{S}_{kt}}{\sum_k p_{kt}}.$$

**Until** convergence **return** $\pi_t, \boldsymbol{U}_t$

### The details of the sampling distributions in the MCMC algorithm

In this section, we dive into the details of the closed forms of the sampling distributions in the MCMC algorithm. We first obtain the sampling distribution of $\boldsymbol{B}_{j\cdot}$. Let $\boldsymbol{e}_j = \widehat{\boldsymbol{B}} - \boldsymbol{R}_{-j}\boldsymbol{B}_{-j\cdot}$ as the residual of the observed eQTL summary statistics ruling out the genetic effects except for SNP $j$, where $\boldsymbol{e}_j|\boldsymbol{B}_{j\cdot} \sim MN(\boldsymbol{R}_j\boldsymbol{B}_{j\cdot}^T, \boldsymbol{R}, \boldsymbol{V})$. Since $\boldsymbol{B}_{j\cdot}|g_j = t \sim \mathcal{N}(\boldsymbol{0}, \boldsymbol{U}_t)$, and $\boldsymbol{R}_j\boldsymbol{B}_{j\cdot}^T \sim MN(\boldsymbol{0}, \boldsymbol{R}_j\boldsymbol{R}_j^T, \boldsymbol{U}_t)$, we have,

$$vec(\boldsymbol{e}_j^T)|g_j = t \sim \mathcal{N}(\boldsymbol{0}, \boldsymbol{R}_j\boldsymbol{R}_j^T \otimes \boldsymbol{U}_t + \boldsymbol{R} \otimes \boldsymbol{V}).$$

In addition,

$$\begin{aligned} Cov(vec(\boldsymbol{e}_j^T), \boldsymbol{B}_{j\cdot}|g_j) &= Cov(vec(\boldsymbol{B}_{j\cdot}\boldsymbol{R}_j^T), \boldsymbol{B}_{j\cdot}|g_j) \\ &= Cov\left((\boldsymbol{R}_j \otimes \boldsymbol{I}_C)\boldsymbol{B}_{j\cdot}, \boldsymbol{B}_{j\cdot}\middle|g_j\right) \\ &= (\boldsymbol{R}_j \otimes \boldsymbol{I}_C)\boldsymbol{U}_t. \end{aligned}$$

Denote $\boldsymbol{T}_{jt} = \boldsymbol{R}_j\boldsymbol{R}_j^T \otimes \boldsymbol{U}_t + \boldsymbol{R} \otimes \boldsymbol{V} \in \mathbb{R}^{MC\times MC}$ . We have

$$\begin{pmatrix} \boldsymbol{B}_{j\cdot} \\ vec(\boldsymbol{e}_j^T) \end{pmatrix} |g_j = t \sim \mathcal{N}\left[\boldsymbol{0}, \begin{pmatrix} \boldsymbol{U}_t & \boldsymbol{U}_t(\boldsymbol{R}_j^T \otimes \boldsymbol{I}_C) \\ (\boldsymbol{R}_j \otimes \boldsymbol{I}_C)\boldsymbol{U}_t & \boldsymbol{T}_{jt} \end{pmatrix}\right],$$

and the conditional distribution

$$\boldsymbol{B}_{j\cdot}|g_j = t, \widehat{\boldsymbol{B}}, \boldsymbol{B}_{-j\cdot} \sim \mathcal{N}\left(\boldsymbol{U}_t(\boldsymbol{R}_j^T \otimes \boldsymbol{I}_C)\boldsymbol{T}_{jt}^{-1}vec(\boldsymbol{e}_j^T), \boldsymbol{U}_t - \boldsymbol{U}_t(\boldsymbol{R}_j^T \otimes \boldsymbol{I}_C)\boldsymbol{T}_{jt}^{-1}(\boldsymbol{R}_j \otimes \boldsymbol{I}_C)\boldsymbol{U}_t\right). \quad (1)$$

Here, $\boldsymbol{B}_{j\cdot}$ is a $C$ vector while the distribution requires calculation in the dimension $MC \times MC$. To reduce the dimensionality to reduce the computation burden, we further simplify the Equation (1).

We first introduce three necessary equations: (1) $(\boldsymbol{A}+\boldsymbol{B})^{-1}=(\boldsymbol{I}+\boldsymbol{A}^{-1}\boldsymbol{B})^{-1}\boldsymbol{A}^{-1}$, which can be easily proved by multiplying $(\boldsymbol{A}+\boldsymbol{B})$ on both sides of the equation; (2) $(\boldsymbol{A}\otimes\boldsymbol{B})(\boldsymbol{C}\otimes\boldsymbol{D})=(\boldsymbol{AC}\otimes\boldsymbol{BD})$; and (3) $(\boldsymbol{A}\otimes\boldsymbol{B})^{-1}=(\boldsymbol{A}^{-1}\otimes\boldsymbol{B}^{-1})$. Then,

$$\boldsymbol{T}_{jt}^{-1}=\left(\boldsymbol{R}_j\boldsymbol{R}_j^T\otimes\boldsymbol{U}_t+\boldsymbol{R}\otimes\boldsymbol{V}\right)^{-1}$$

$$=\left(\boldsymbol{I}+\boldsymbol{R}^{-1}\boldsymbol{R}_j\boldsymbol{R}_j^T\otimes\boldsymbol{V}^{-1}\boldsymbol{U}_t\right)^{-1}(\boldsymbol{R}^{-1}\otimes\boldsymbol{V}^{-1})$$

$$=\left(\boldsymbol{I}+\boldsymbol{l}_j\boldsymbol{R}_j^T\otimes\boldsymbol{V}^{-1}\boldsymbol{U}_t\right)^{-1}(\boldsymbol{R}^{-1}\otimes\boldsymbol{V}^{-1}).$$

Here, $\boldsymbol{l}_j$ is an $M$-vector containing all zeros except for a one in the $j$th position, $\boldsymbol{R}_j^T\boldsymbol{l}_j=R_{jj}=1$ and $\boldsymbol{l}_j\boldsymbol{R}_j^T=\boldsymbol{l}_j\otimes\boldsymbol{R}_j^T$. Thus, $\left(\boldsymbol{I}+\boldsymbol{l}_j\boldsymbol{R}_j^T\otimes\boldsymbol{V}^{-1}\boldsymbol{U}_t\right)\boldsymbol{T}_{jt}^{-1}=\boldsymbol{R}^{-1}\otimes\boldsymbol{V}^{-1}$, i.e., $\boldsymbol{T}_{jt}^{-1}=\boldsymbol{R}^{-1}\otimes\boldsymbol{V}^{-1}-\boldsymbol{l}_j\otimes\left[\left(\boldsymbol{R}_j^T\otimes\boldsymbol{V}^{-1}\boldsymbol{U}_t\right)\boldsymbol{T}_{jt}^{-1}\right]$. Denote $\boldsymbol{A}=\left(\boldsymbol{R}_j^T\otimes\boldsymbol{V}^{-1}\boldsymbol{U}_t\right)\boldsymbol{T}_{jt}^{-1}$. Then,

$$\left(\boldsymbol{I}+\boldsymbol{l}_j\boldsymbol{R}_j^T\otimes\boldsymbol{V}^{-1}\boldsymbol{U}_t\right)\left(\boldsymbol{R}^{-1}\otimes\boldsymbol{V}^{-1}-\boldsymbol{l}_j\otimes\boldsymbol{A}\right)=\boldsymbol{R}^{-1}\otimes\boldsymbol{V}^{-1}$$

$$\Leftrightarrow\boldsymbol{l}_j\boldsymbol{R}_j^T\boldsymbol{R}^{-1}\otimes\boldsymbol{V}^{-1}\boldsymbol{U}_t\boldsymbol{V}^{-1}=\boldsymbol{l}_j\otimes\boldsymbol{A}+\boldsymbol{l}_j\boldsymbol{R}_j^T\boldsymbol{l}_j\otimes\boldsymbol{V}^{-1}\boldsymbol{U}_t\boldsymbol{A}$$

$$\Leftrightarrow\boldsymbol{R}_j^T\boldsymbol{R}^{-1}\otimes\boldsymbol{V}^{-1}\boldsymbol{U}_t\boldsymbol{V}^{-1}=\boldsymbol{A}+\boldsymbol{V}^{-1}\boldsymbol{U}_t\boldsymbol{A}$$

$$\Leftrightarrow\boldsymbol{A}=(\boldsymbol{I}+\boldsymbol{V}^{-1}\boldsymbol{U}_t)^{-1}\left(\boldsymbol{l}_j^T\otimes\boldsymbol{V}^{-1}\boldsymbol{U}_t\boldsymbol{V}^{-1}\right)$$

$$\Leftrightarrow\boldsymbol{A}=\boldsymbol{l}_j^T\otimes[(\boldsymbol{I}+\boldsymbol{V}^{-1}\boldsymbol{U}_t)^{-1}\boldsymbol{V}^{-1}\boldsymbol{U}_t\boldsymbol{V}^{-1}]$$

$$\Leftrightarrow\boldsymbol{A}=\boldsymbol{l}_j^T\otimes[(\boldsymbol{U}_t+\boldsymbol{V})^{-1}\boldsymbol{U}_t\boldsymbol{V}^{-1}]$$

$$\Leftrightarrow\boldsymbol{A}=\boldsymbol{l}_j^T\otimes[\boldsymbol{V}^{-1}-(\boldsymbol{U}_t+\boldsymbol{V})^{-1}].$$

Therefore,

$$\boldsymbol{T}_{jt}^{-1}=\boldsymbol{R}^{-1}\otimes\boldsymbol{V}^{-1}-\boldsymbol{l}_j\boldsymbol{l}_j^T\otimes[\boldsymbol{V}^{-1}-(\boldsymbol{U}_t+\boldsymbol{V})^{-1}],$$

$$\left(\boldsymbol{R}_j^T\otimes\boldsymbol{I}_C\right)\boldsymbol{T}_{jt}^{-1}=\boldsymbol{R}_j^T\boldsymbol{R}^{-1}\otimes\boldsymbol{V}^{-1}-\boldsymbol{R}_j^T\boldsymbol{l}_j\boldsymbol{l}_j^T\otimes[\boldsymbol{V}^{-1}-(\boldsymbol{U}_t+\boldsymbol{V})^{-1}]=\boldsymbol{l}_j^T\otimes(\boldsymbol{U}_t+\boldsymbol{V})^{-1},$$

$$\left(\boldsymbol{R}_j^T\otimes\boldsymbol{I}_C\right)\boldsymbol{T}_{jt}^{-1}\left(\boldsymbol{R}_j\otimes\boldsymbol{I}_C\right)=\boldsymbol{l}_j^T\boldsymbol{R}_j\otimes(\boldsymbol{U}_t+\boldsymbol{V})^{-1}=(\boldsymbol{U}_t+\boldsymbol{V})^{-1}.$$

The conditional mean in Equation (1) is

$$\boldsymbol{U}_t\left(\boldsymbol{R}_j^T \otimes \boldsymbol{I}_C\right)\boldsymbol{T}_{jt}^{-1} vec\left(\boldsymbol{e}_j^T\right) = \boldsymbol{U}_t\left[\boldsymbol{l}_j^T \otimes (\boldsymbol{U}_t + \boldsymbol{V})^{-1}\right] vec\left(\boldsymbol{e}_j^T\right) = \boldsymbol{U}_t(\boldsymbol{U}_t + \boldsymbol{V})^{-1}\boldsymbol{e}_{j,j},$$

where $\boldsymbol{e}_{j,j} = \widehat{\boldsymbol{B}}_{j\cdot} - \boldsymbol{R}_{j,-j}^T \boldsymbol{B}_{-j\cdot}$ is the $j$th row of $\boldsymbol{e}_j$. The conditional variance in Equation (1) is

$$\boldsymbol{U}_t - \boldsymbol{U}_t\left(\boldsymbol{R}_j^T \otimes \boldsymbol{I}_C\right)\boldsymbol{T}_{jt}^{-1}\left(\boldsymbol{R}_j \otimes \boldsymbol{I}_C\right)\boldsymbol{U}_t = \boldsymbol{U}_t - \boldsymbol{U}_t(\boldsymbol{U}_t + \boldsymbol{V})^{-1}\boldsymbol{U}_t.$$

Thus,

$$\boldsymbol{B}_{j\cdot}|g_j = t, \widehat{\boldsymbol{B}}, \boldsymbol{B}_{-j\cdot} \sim \mathcal{N}\left(\boldsymbol{U}_t(\boldsymbol{U}_t + \boldsymbol{V})^{-1}\boldsymbol{e}_{j,j}, \boldsymbol{U}_t - \boldsymbol{U}_t(\boldsymbol{U}_t + \boldsymbol{V})^{-1}\boldsymbol{U}_t\right). \quad (2)$$

Next, we obtain the sampling distribution for $g_j$:

$$P\left(g_j = t \middle| vec\left(\boldsymbol{e}_j^T\right)\right) = \frac{\pi_t f\left(vec\left(\boldsymbol{e}_j^T\right)|g_j = t\right)}{\sum_{s=1}^T \pi_s f\left(vec\left(\boldsymbol{e}_j^T\right)|g_j = s\right)}. \quad (3)$$

The following equalities are all under the equivalence for any constant unrelated of $t$. Since $vec\left(\boldsymbol{e}_j^T\right)|g_j = t$ follows a normal distribution, we have

$$\ln f\left(vec\left(\boldsymbol{e}_j^T\right)|g_j = t\right) = \frac{1}{2}\ln\det \boldsymbol{T}_{jt}^{-1} - \frac{1}{2} vec\left(\boldsymbol{e}_j^T\right)^T \boldsymbol{T}_{jt}^{-1} vec\left(\boldsymbol{e}_j^T\right).$$

For the first term,

$$\begin{aligned}\ln\det \boldsymbol{T}_{jt}^{-1} &= \ln\det\left(\boldsymbol{R}^{-1} \otimes \boldsymbol{V}^{-1} - \boldsymbol{l}_j\boldsymbol{l}_j^T \otimes [\boldsymbol{V}^{-1} - (\boldsymbol{U}_t + \boldsymbol{V})^{-1}]\right) \\ &= \ln\det(\boldsymbol{R}^{-1} \otimes \boldsymbol{V}^{-1}) + \ln\det\left(\boldsymbol{I} + \boldsymbol{R}\boldsymbol{l}_j\boldsymbol{l}_j^T \otimes [\boldsymbol{V}(\boldsymbol{U}_t + \boldsymbol{V})^{-1} - \boldsymbol{I}]\right) \\ &= \ln\det(\boldsymbol{V}(\boldsymbol{U}_t + \boldsymbol{V})^{-1}) \\ &= -\ln\det(\boldsymbol{U}_t + \boldsymbol{V}).\end{aligned}$$

For the second term,

$$\begin{aligned} vec\left(\boldsymbol{e}_j^T\right)\boldsymbol{T}_{jt}^{-1} vec\left(\boldsymbol{e}_j^T\right) &= -vec\left(\boldsymbol{e}_j^T\right)^T\left(\boldsymbol{l}_j\boldsymbol{l}_j^T \otimes [\boldsymbol{V}^{-1} - (\boldsymbol{U}_t + \boldsymbol{V})^{-1}]\right) vec\left(\boldsymbol{e}_j^T\right) \\ &= +\boldsymbol{e}_{j,j}^T[(\boldsymbol{U}_t + \boldsymbol{V})^{-1} - \boldsymbol{V}^{-1}]\boldsymbol{e}_{j,j}.\end{aligned}$$

Thus,

$$\ln f\left(vec\left(\boldsymbol{e}_j^T\right)\middle|g_j = t\right) = -\frac{1}{2}\ln\det(\boldsymbol{U}_t + \boldsymbol{V}) - \frac{1}{2}\boldsymbol{e}_{j,j}^T[(\boldsymbol{U}_t + \boldsymbol{V})^{-1} - \boldsymbol{V}^{-1}]\boldsymbol{e}_{j,j}.$$

In conclusion, for each SNP $j$, we first use Equation (3) to first sample a category $g_j$, and then use Equation (2) to sample the true effect sizes $\boldsymbol{B}_{j\cdot}$.

<u>**Algorithm 2** MCMC: Sampling from the posterior distributions**:**</u>

**Require:** eQTL summary statistics, $\widehat{\boldsymbol{B}}$, LD matrix $\boldsymbol{R}$, sample-adjusted covariance matrix $\widehat{\boldsymbol{V}}$, prior parameters, $\pi_t, \boldsymbol{U}_t$.

Initialize eQTL effect size matrices, $\boldsymbol{B}^{(0)}$, and the hidden indicators, $\boldsymbol{g}^{(0)}$.

**Repeat**

For $k$ in $1, \cdots, M$ do

$\boldsymbol{e}_k = \widehat{\boldsymbol{B}}_k - \boldsymbol{R}_{k,-k}\boldsymbol{B}_{-k}^{(w)}$.

$\boldsymbol{S}_t = (\boldsymbol{U}_t + \boldsymbol{V})^{-1}$.

$p_t = P\left(g_k = t\middle|vec(\boldsymbol{e}_k^T)\right) \propto \pi_t \det(\boldsymbol{S}_t)^{\frac{1}{2}} \exp(\boldsymbol{e}_k^T\boldsymbol{S}_t\boldsymbol{e}_k)^{\frac{1}{2}}$.

Randomly sample $g_k^{(w+1)} \sim Multinom(\boldsymbol{p})$.

Randomly sample $\boldsymbol{B}_k^{(w+1)} \sim \mathcal{N}(\boldsymbol{U}_t\boldsymbol{S}_t\boldsymbol{e}_k, \boldsymbol{U}_t - \boldsymbol{U}_t\boldsymbol{S}_t\boldsymbol{U}_t)$, for $t = g_k^{(w+1)}$.

**End for**

**Until** convergence **return** $\boldsymbol{g}^{(w)}, \boldsymbol{B}^{(w)}, w = 1, \cdots W$.

### Supplementary Figures

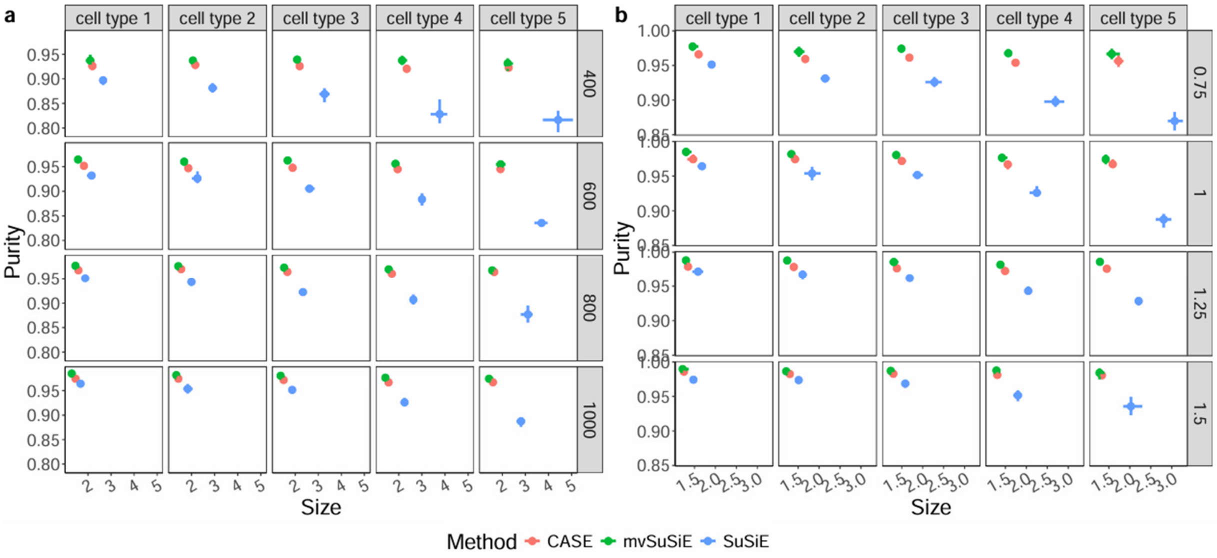


**Supplementary Fig. 1. Examples of Fine-mapping results.** We compare CASE (red), mvSuSiE (green), and SuSiE (blue) in simulation studies for the Purity and Size in varied sample size (**a**) and varied heritability multiplier (**b**). Each block represents a cell type indexed from one to five in a simulation setting. Error bars indicate upper 90% and lower 10% percentiles across 100 simulations.

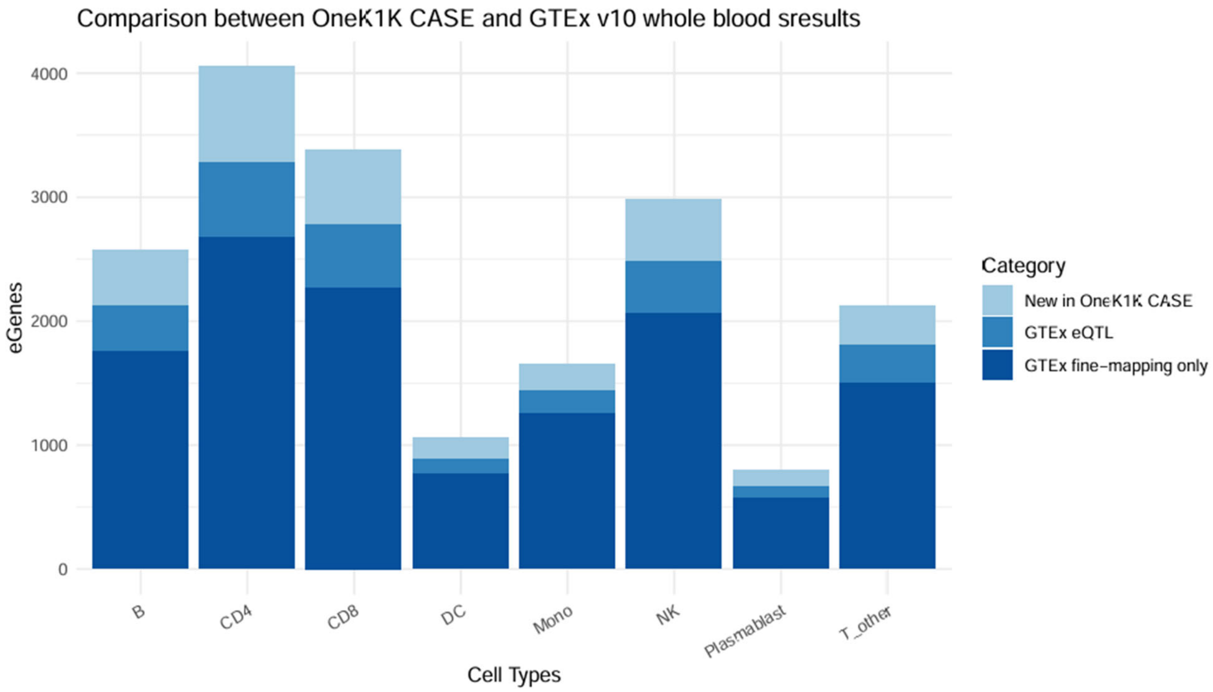


**Supplementary Fig. 2. Comparison between CASE OneK1K results and GTEx v10 whole blood results.** We compare the eGenes across cell types. The dark blue represents those eGenes only found by GTEx fine-mapping results, the medium blue represents additional eGenes found by GTEx eQTL marginal associations, and the light blue represents the novel ones identified in CASE OneK1K application.

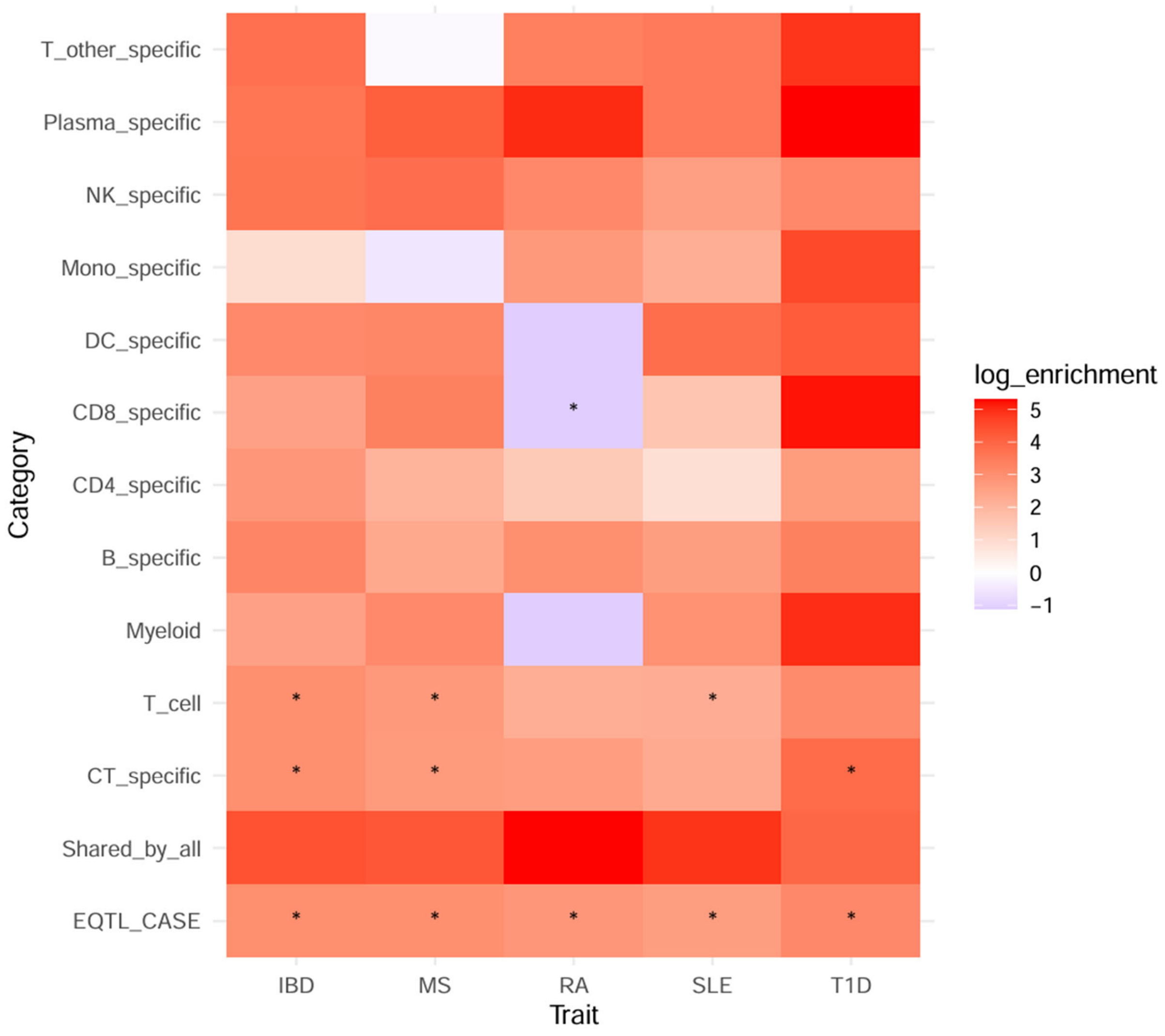


**Supplementary Fig. 3. Heritability enrichment analyses for CASE-identified eQTLs.** The x-axis represents diseases (*IBD*: Inflammatory Bowel Disease; *MS*: Multiple Sclerosis; *RA*: Rheumatoid Arthritis; *SLE*: Systemic Lupus Erythematosus; *T1D*: Type 1 Diabetes) and the y-axis represents different categories of CASE-identified eQTLs, including all the eQTLs (*EQTL_CASE*), universally shared eQTLs (*Shared_by_all*), eQTLs unique to one cell type (*CT_specific*), eQTLs only shared within T cells (*T_cell*), eQTLs only shared within myeloid cells (*Myeloid*), and cell-type-specific eQTLs for each cell type. We use asterisks for significant disease-category pairs (adjusted p-values $\leq 0.1$).

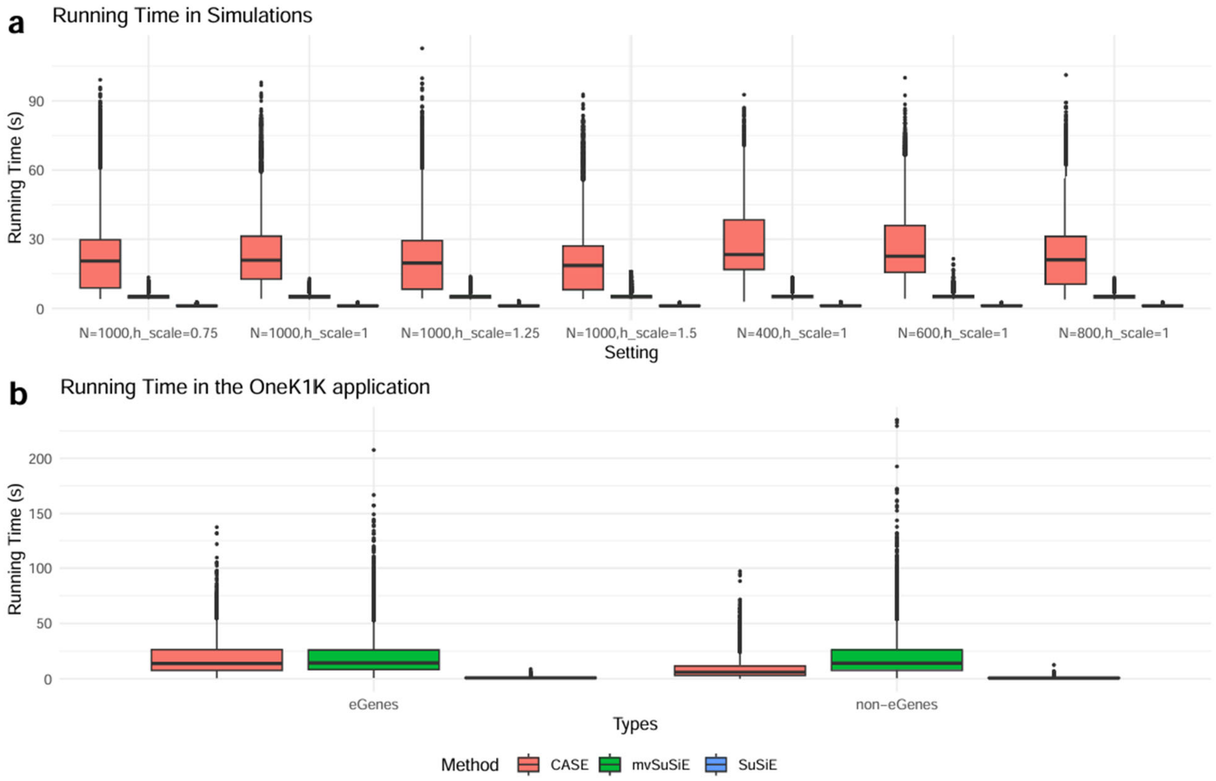


**Supplementary Fig. 4. Running time of CASE, mvSuSiE and SuSiE in simulations and real applications.** We compare the running time of CASE (red), mvSuSiE (green), and SuSiE (blue) in simulation studies (**a**) and real applications (**b**). **a.** the x-axis represents each simulation setting with respect to the sample size (N) and heritability multiplier (h_scale). **b.** the x-axis represents scenarios whether the genes are identified as eGenes by CASE.

### Supplementary Table 1. Information of Cell Types in the OneK1K Dataset

| Cell type Abbreviations | Subtypes | # of Cells | | Sample size |
|---|---|---|---|---|
| CD4 | CD4-positive, alpha-beta T cell | 773 | 598,039 | 981 |
| | naive thymus-derived CD4-positive, alpha-beta T cell | 259,012 | | |
| | central memory CD4-positive, alpha-beta T cell | 289,000 | | |
| | CD4-positive, alpha-beta cytotoxic T cell | 17,993 | | |
| | effector memory CD4-positive, alpha-beta T cell | 31,261 | | |
| CD8 | effector memory CD8-positive, alpha-beta T cell | 161,051 | 230,303 | 981 |
| | central memory CD8-positive, alpha-beta T cell | 16,409 | | |
| | naive thymus-derived CD8-positive, alpha-beta T cell | 52,538 | | |
| | CD8-positive, alpha-beta T cell | 305 | | |
| NK | natural killer cell | 164,933 | 171,939 | 981 |
| | CD16-negative, CD56-bright natural killer cell | 7,006 | | |
| B | memory B cell | 30,234 | 125,825 | 981 |
| | naive B cell | 65,702 | | |
| | transitional stage B cell | 29,889 | | |
| T_other | gamma-delta T cell | 18,922 | 56,112 | 981 |
| | regulatory T cell | 26,531 | | |
| | mucosal invariant T cell | 8,835 | | |
| Mono | CD14-positive monocyte | 36,130 | 51,873 | 978 |
| | CD14-low, CD16-positive monocyte | 15,743 | | |
| DC | dendritic cell | 181 | 6,648 | 921 |
| | plasmacytoid dendritic cell | 1,897 | | |
| | conventional dendritic cell | 4,570 | | |
| Plasma | plasma blast | 3,754 | 3,754 | 795 |
| [Removed] | double negative thymocyte | 1,824 | / | / |
| | peripheral blood mononuclear cell | 131 | | |
| | hematopoietic precursor cell | 1,812 | | |
| | erythrocyte | 290 | | |
| | platelet | 1,810 | | |
| | innate lymphoid cell | 444 | | |
